# Unscheduled origin building in S-phase upon tight CDK1 inhibition suppresses CFS instability

**DOI:** 10.1101/2020.11.19.390054

**Authors:** Olivier Brison, Stefano Gnan, Dana Azar, Mélanie Schmidt, Stéphane Koundrioukoff, Sami El-Hilali, Yan Jaszczyszyn, Anne-Marie Lachages, Claude Thermes, Chun-Long Chen, Michelle Debatisse

## Abstract

Genome integrity requires replication to be completed before chromosome segregation. This coordination essentially relies on replication-dependent activation of a dedicated checkpoint that inhibits CDK1, delaying mitotic onset. Under-replication of Common Fragile Sites (CFSs) however escapes surveillance, which triggers chromosome breakage. Using human cells, we asked here whether such leakage results from insufficient CDK1 inhibition under modest stresses used to destabilize CFSs. We found that tight CDK1 inhibition suppresses CFS instability. Repli-Seq and molecular combing analyses consistently showed a burst of replication initiations in mid S phase across large origin-poor domains shaped by transcription, including CFSs. Strikingly, CDC6 or CDT1 depletion or CDC7-DBF4 inhibition during the S phase prevented both extra-initiations and CFS rescue, showing that CDK1 inhibition promotes targeted and mistimed building of functional extra-origins. In addition to delay mitotic onset, checkpoint activation therefore advances replication completion of chromosome domains at risk of under-replication, two complementary roles preserving genome stability.

## Introduction

At each cell cycle, the replication process should duplicate the entire genome, in normal conditions as well as under replication stress. Licensing of tens of thousands of loci to become replication origins contributes to solve the challenge raised by the completion of human genome duplication within the time frame of the S phase ^1^. Licensing starts with chromatin loading of the ORC complex and CDC6 that serve as a platform for sequential CDT1-driven recruitment of two head-to-head MCM hexamers, constituting the pre-Replication Complex (pre-RC) ^2–5^. Pre-RC building starts in late mitosis and continues during the G1 phase. At the onset of the S phase, phosphorylation events by cyclin-dependent kinases (CDKs) and the DBF4-dependent kinase (CDC7-DBF4) elicit the binding of additional factors to form the CMG (Cdc45-MCM-GINS) complex, the functional replicative helicase. A third round of protein recruitment and post-translational modifications then timely converts the two CMG complexes into an active origin from which bi-directional replication forks emanate ^6,7^. It is now well established that only a subset of the pre-RCs generates replication forks in a normal cell cycle and that supernumerary pre-RCs provide a pool of salvage origins allowing the cells to increase the density of initiation events when required. This adaptation process, called compensation, helps replication to proceed when cells undergo replication stress ^8^.

In addition to compensation, eukaryotic cells have evolved processes that monitor S-phase progression and prevent mitotic entry before replication completion in normal conditions as well as under exogenous replication stress ^9^. It has long been demonstrated that severe replication stresses activate ATR, the apical kinase of the DNA Replication Checkpoint (DRC), and downstream CHK1 kinase, which triggers complex phosphorylation cascades. A major outcome of the pathway is inhibition of CDK1, the activity of which is mandatory for mitotic entry. Recent reports have now shown that DNA replication *per se* elicits a signal restraining mitotic onset in cells undergoing a normal S phase. Indeed, ongoing replication forks activate the DRC ^10, 11^, which delays CDK1-dependent activation of FOXM1 ^11^, an important trans-activator of the mitotic transcription program ^12^. An ATR/CHK1-independent attenuation of CDK1 activity has also been observed in cells challenged by modest replication stress. This attenuation results from sequestration of CDK1 in specific nuclear foci, an environment in which its kinase activity is reduced ^13^. Various mechanisms therefore modulate CDK1 activity in order to coordinate replication completion to mitotic entry, both in cells submitted to exogenous stress and in cells undergoing only endogenous stress.

A prominent source of endogenous replication stress and ensuing genome instability is transcription that perturbs replication by two major mechanisms; first, through interference between the replication and the transcription machines, notably head-on encounters that favor R-loop stabilization and subsequent replication fork stalling ^14–16^. This mechanism has notably been involved in the instability of early-replicating fragile sites (ERFSs), a category of hard-to-replicate regions associated with small genes early replicating and highly expressed ^17–19^; second through transcription-dependent segregation of initiation events out of expressed genes ^20–26^. This reorganization of the replication initiation profiles tends to co-orientate replication and transcription, minimizing head-on encounters. In addition, when co-oriented with transcription, replication forks tend to clear spontaneously formed RNA-DNA hybrids, alleviating their potential toxicity ^27^. While globally beneficial, initiation paucity along the gene body becomes highly detrimental for large genes, the replication of which consequently relies on long-traveling forks emanating from their flanking regions. Stress-induced fork slowing therefore specifically delays replication completion of these genes, explaining the vulnerability of common fragile sites (CFSs), another category of hard-to-replicate regions nested in large genes late replicating and moderately expressed ^28, 29^.

CFS instability increases in various genetic contexts, notably ATR or CHK1 deficiency ^30, 31^. It remains however unclear why primary lymphocytes and fibroblasts from individuals with proficient DRC breach surveillance to enter mitosis with incompletely replicated CFSs ^28^. A possible explanation relies on the observation that lymphoblasts challenged with low doses of aphidicolin (Aph), an inhibitor of replicative DNA polymerases classically used to destabilize CFSs ^32^, display recruitment of ATR on the chromatin but weak CHK1, p53 and γ-H2AX phosphorylation ^33^. Hence, CDK1 inhibition may be insufficient to ensure tight control on mitotic onset in so-treated cells. Here we showed that Aph-induced CFS instability is completely suppressed in human lymphoblasts and fibroblast upon co-treatment of the cells with RO3306 (RO), a reversible CDK1 inhibitor ^34^. We also treated lymphoblasts with low doses of hydroxyurea (HU), a potent ribonucleotide reductase inhibitor. In contrast to Aph, low doses of HU efficiently activate CHK1 and downstream effectors. This difference most probably results from the impact of HU on precursor pools. Indeed, we have previously shown in JEFF cells that low HU concentrations strongly reduce the dATP pool and increase the dTTP pool ^35^. Resulting pool unbalance may enhance mutagenesis and challenge genome integrity by mechanisms largely independent of fork slowing *per se* ^36^. Cytogenetic analysis of HU-treated cells showed here that CFSs are far less instable in cells treated with HU than in cells treated with Aph for similar levels of fork slowing, strongly supporting the idea that tight DRC activation rescues CFSs.

To decipher the mechanisms responsible for this rescue, we compared the replication dynamics in lymphoblasts treated with Aph to that of cells treated with HU. We also treated the cells with Aph plus RO (ARO) or, conversely, with HU plus the ATR inhibitor VE821 (HU+ATRi) to prevent DRC-induced CDK1 inhibition. Repli-Seq analyses revealed, specifically in cells in which CDK1 is inactive, a burst of neo-synthesized DNA along large genes, including those hosting CFSs, that starts in mid S phase and results in strong advancement of their replication time. Focusing on a canonical CFS, we showed by molecular combing that extra initiation events account for this burst of DNA synthesis. We further showed that availability, during the S phase, of proteins involved in origin setting is mandatory for both rescue of CFS stability and extra-initiations in cells treated with ARO. Together, results show that DRC-induced CDK1 inhibition promotes mistimed building of new pre-RCs, then of functional origins. Firing of these origins accelerates replication completion of transcribed large genes and some other types of late-replicating origin-poor domains, without promoting global re-replication. One prominent role of DRC activation is therefore to advance the replication timing of genomic regions at risk of under-replication. This yet unsuspected checkpoint outcome most probably plays a major role in the prevention of CFS-induced chromosomal rearrangements driving oncogenesis ^37^ and human diseases, notably neurodegenerative and neuropsychiatric diseases ^38^.

## Results

### RO reversibly blocks cells in G2 phase without triggering DNA damage

We used RO to reversibly inhibit CDK1 (Figure 1A), a drug shown to be 10 times more specific of CDK1 (Ki=35 nM) than of CDK2, the nearest related kinase (Ki=340 nM) (Vassilev, 2006). At 10 μM, RO permits most JEFF lymphoblasts (human B-lymphocytes immortalized by Epstein-Barr virus) to accumulate with a 4C DNA content after 16 h of treatment, *i.e*. approximately one doubling time (Figures 1B, cell cycle t16/0 h and S1A). These cells were blocked in G2 phase since very few mitotic cells were present in the population (Figure 1B, mitotic cells t16/0 h). Comparable cell accumulation was also observed with MRC5 human primary fibroblasts (Figure S1B) and GMA32 immortalized Chinese hamster fibroblasts (Figure S1C). These results confirm previous works ^34, 39, 40^ showing that most cells enter and complete S phase in the absence of CDK1 activity, and stop cycling in G2 phase.

**Figure 1:**
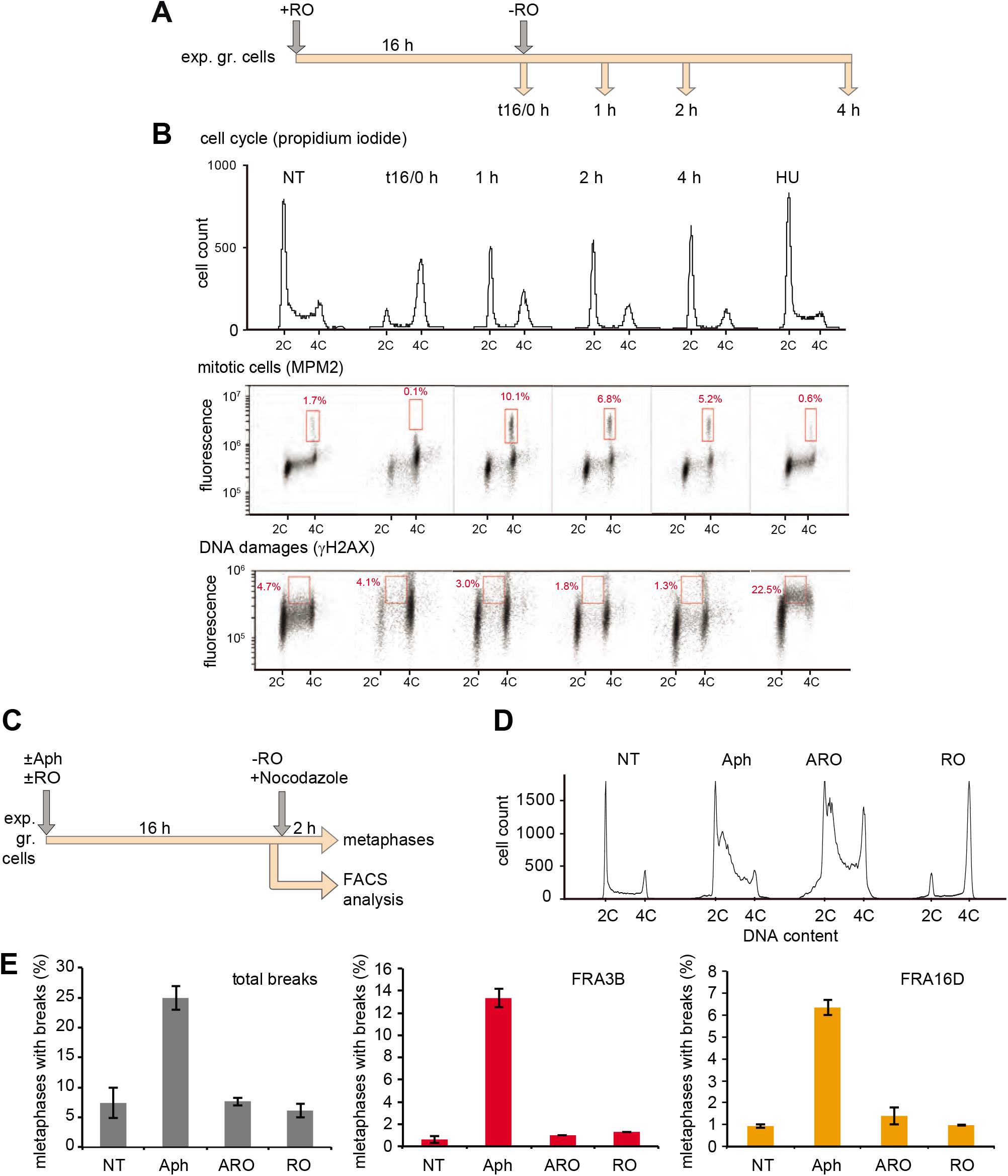
RO reversibly blocks cells in G2 phase without triggering DNA damage, and suppresses Aph-induced CFS instability. **A:** Experimental scheme. Exponentially growing (exp. gr.) JEFF cells were incubated for 16 h in the presence of RO (10 μM). Cells were then released in normal medium (t16/0 h) and samples were collected at indicated times. **B:** Cells treated as described in A were stained with propidium iodine and analyzed by FACS (cell cycle) or labeled with antibodies anti-MPM2 (mitotic cells) or anti-γH2AX (DNA damages). Percentages of mitotic cells or γH2AX positive cells (red squares) are indicated. Non-treated cells (NT) or cells treated for 2 h with HU 1 mM (HU) were included as controls. **C:** Experimental scheme. Exponentially growing cells were incubated for 16 h in the presence or absence of Aph (600 nM) and/or RO (10 μM). **D:** Part of the cells was fixed for FACS analysis. **E:** After RO removal the rest of the cells was treated with nocodazole (2 h, 200 nM) and metaphases were collected. The frequency of total chromosome breaks was determined on Giemsa stained chromosomes (Figure S1G) and breaks at FRA3B or FRA16D were scored after DAPI staining and FISH with probes specific to the *FHIT* or *WWOX* gene (Figure S1H). Experiments shown in E were carried out twice and error bars represent standard deviation.

We also found that JEFF cells accumulated in G2 phase after 16 h of RO-treatment do not display γ-H2AX signal (Figures 1B, DNA damages t16/0 h and S1D). When RO was washed out at that time, a majority of G2 arrested cells entered and completed mitosis (Figure 1B, cell cycle and mitotic cells). Cytogenetic analyses showed that the percentage of Giemsa-stained metaphase plates displaying breaks on any chromosome (total breaks) does not significantly increase after release relative to untreated cells (Figure 1E, RO/NT). In addition, JEFF cells that reached and progressed in the following G1 phase up to S phase displayed no γ-H2AX accumulation (Figures 1B, DNA damages and S1D). Consistent results were obtained with GMA32 cells (Figures S1E and S1F). Therefore, the absence of CDK1 activity up to one cell cycle does not induce DNA damages, this feature being neither species-nor cell type-specific.

### RO suppresses Aph-induced CFS instability without reducing large gene transcription

To determine how RO impacts cell cycle progression and chromosome stability in cells submitted to modest replication stress, JEFF cells were incubated for 16 h in the presence of Aph 600 nM, a concentration routinely used to induce breaks at CFSs, or RO 10 μM, or ARO (Aph 600 nM and RO 10 μM) (Figure 1C). Unless specifically mentioned, these concentrations were used in all subsequent experiments. FACS analyses showed that JEFF cells treated with Aph tend to accumulate in early S phase. This feature was also observed upon growth in ARO although another part of the cells accumulates in G2 phase (Figure 1D). Therefore, RO does not interfere with Aph-dependent perturbation of S phase progression and Aph does not interfere with RO-dependent bloc to mitotic onset. The percentages of total breaks were determined in the four growth conditions (Figures 1E and S1G), and fluorescence in situ hybridization (FISH) was carried out to assess the fragility of FRA3B and FRA16D (Figure S1H), two major CFSs in primary lymphocytes ^32^ and in JEFF cells ^29^. Total breaks were also analyzed in MRC5 cells (Figure S1G), as well as breaks at FRA1L and FRA3L (Figure S1H), the two major CFSs in these cells ^41^. We found that RO completely suppresses Aph-induced total breaks and, consistently, breaks at CFSs in either cell type (Figures 1E and S1I).

Since it is now well established that transcription of late replicating large genes dictates CFS setting ^28^, we asked whether the transcriptional status of the *FHIT* (FRA3B) and *WWOX* (FRA16D) genes are impacted in JEFF cells treated for 16 h with Aph, RO or ARO. To this aim, we quantified nascent RNAs by RT-qPCR using a series of intronic primers (Figure S1J). In addition, we performed chromatin immunoprecipitation (ChIP) of RNA polymerase II and qPCR quantification with the same primers as above (Figure S1K). Results showed that none of these treatments significantly reduces the transcription level of the *FHIT* and *WWOX* genes. Similar results were obtained upon study of the *NEGR1* gene (FRA1L) in MRC5 cells (Figure S1J). Thus, rescue of CFSs in cells grown in ARO does not result from attenuated transcription of hosting genes.

### RO favors completion of CFS replication

We used the Repli-Seq technique to compare the effects of a 16 h treatment with Aph or
ARO on genome-wide replication timing profiles. Repli-Seq has previously allowed us to determine how 16 h of treatment with Aph 600 nM impacts the replication dynamics of JEFF cells ^29^. We notably computed an under-replication index (URI) that reflects the difference between the sums of reads per 50 kb window along the genome of cells treated with Aph and untreated cells (Δ_Aph-NT_). Windows with URI≤-2 were recorded as Significantly Delayed/under-replicated Windows (SDWs). The clustering of SDWs identified regions, named Significantly Delayed Regions (SDRs), displaying at least two SDWs separated by less than 250 kb, a distance far smaller than expected from random distribution (P < 10^−15^). We found that over 80 % of the SDRs, called T-SDRs, nest in chromosome domains transcribed continuously across at least 300 kb. Strikingly, SDRs turned to be the landmark of CFSs. Here we compared the replication profile of JEFF cells submitted to various treatments to that of untreated cells (Figure 2A). Briefly, cells pulse-labeled with BrdU were FACS sorted into six fractions (G1/S1, S2 to S5, S6/G2/M) (Figure S2A), newly synthesized DNA was isolated from each fraction, sequenced and the reads were mapped on the human genome (Hg19). After normalization (Figures S2B and S2C) (Methods) data obtained from untreated cells and cells grown in Aph or ARO showed that biological replicates give highly reproducible profiles (R>0.79, P<2.2 10^−16^) (Figures S2D and S2E).

**Figure 2:**
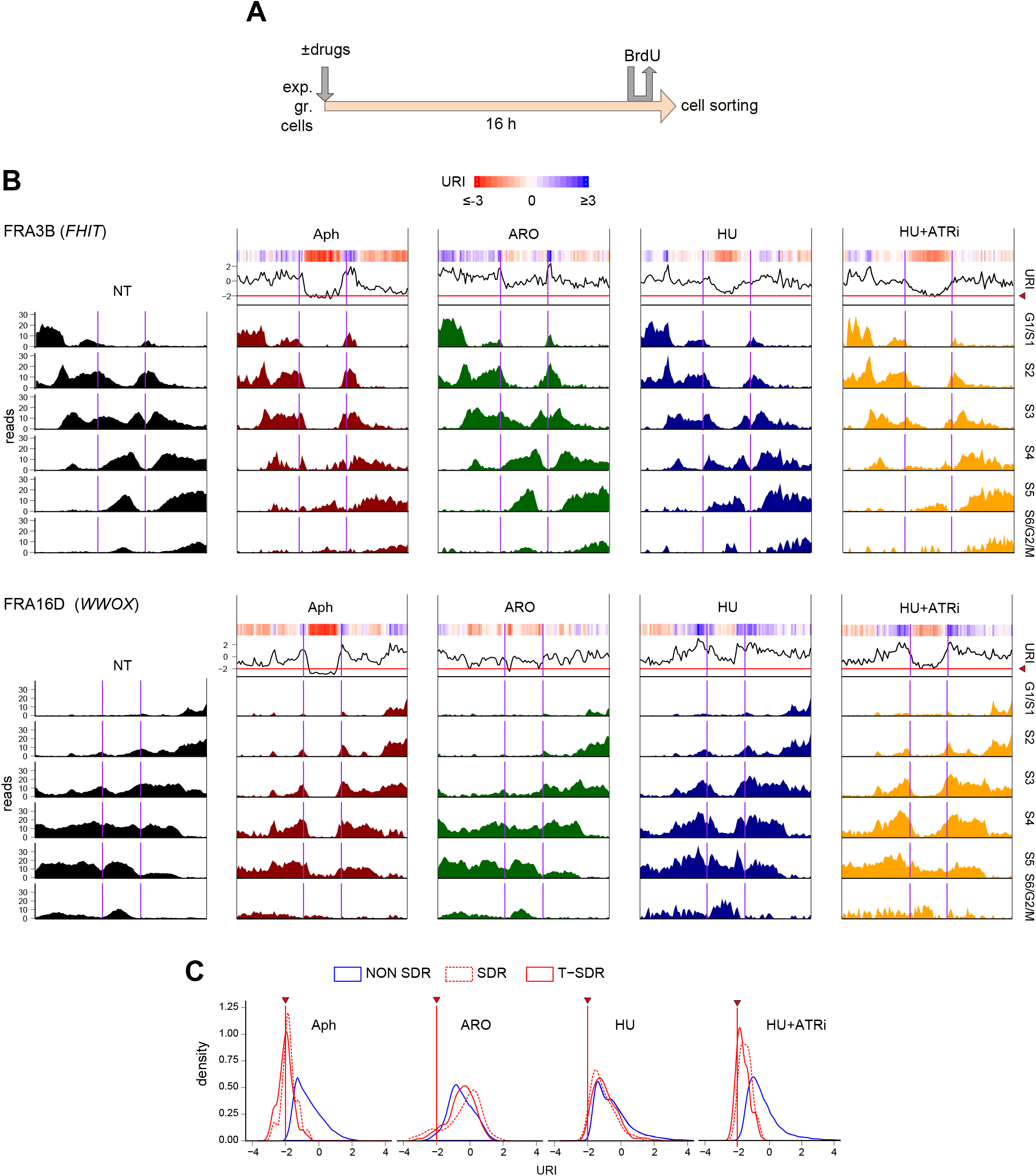
Repli-Seq analyses of URIs along SDR-containing domains in cells treated with Aph, ARO, HU or HU+ATRi. **A:** A schematic representation of the experimental procedure. **B:** Top, heatmap color bar for URIs. Bins with values lower than −3 or higher than 3 are saturated for visualization purposes. Below, mean Repli-Seq profiles at 6 successive stages of the cell cycle (Figure S2A and Methods) along domains overlapping FHIT or WWOX and sequences 2 Mb-long flanking the 5’ and 3’ ends of either gene. Two purples vertical lines delineate the FHIT and WWOX genes, respectively. The same genomic regions with coordinates can be found in figure S2E. The profiles of non-treated cells (NT, black) and cells treated with Aph (red), ARO (green), HU (blue) or HU+ATRi (yellow) are shown. The URIs calculated at 50 kb resolution (ΔStress-NT, Methods) are shown on top of the Repli-Seq data (heatmap and black curve). The horizontal red line highlighted by a red arrowhead crossing the curves of URI profiles indicates the −2 cut-off, i.e. the threshold of significance for delayed replication ^29^. **C:** URI density distribution over 50 kb windows corresponding to SDRs not associated with transcripts (red dashed curve), T-SDRs (red solid curve) and the bulk late replicating regions (NON-SDR, blue curve). Red vertical line with arrowhead: as in B.

We then calculated URIs by computing data from the mean of 2 (Aph and ARO) or 3 (untreated) independent experiments (Methods). Analysis of the profiles obtained along chromosome domains containing the *FHIT* or the *WWOX* genes (Figures 2B and S2E) showed that replication of the genes starts from their flanking regions in G1/S1 or S2, respectively. In untreated cells, replication completion of either gene took place in S6/G2/M. In agreement with our previous results ^29^, we found here with additional independent experiments that cells treated for 16 h with Aph display a large T-SDR in the *FHIT* and *WWOX* genes. In contrast, cells grown in ARO showed high amounts of newly synthesized DNA across the two genes, suppressing the T-SDRs. This replication stimulation strikingly starts from mid S phase, which permits completion of *FHIT* and *WWOX* replication in late S phase (Figures 2B and S2E). Further analyses showed that stimulation of replication in ARO-treated cells concerns all SDRs and T-SDRs, resulting in drastic reduction of windows with URI≤-2 (Figures 2C, 3 and S2F). These results indicate that inhibition of CDK1 by RO stimulates replication of CFSs, which explains their stability in cells grown in ARO, and also exclude the recently proposed hypothesis that G2 phase lengthening passively permits completion of CFS replication ^42^.

### RO stimulates replication of large genes that complete replication past mid-S in Aph

We also determined how Aph impacts the replication dynamics of the bulk genes. We classified genes according to their size, which confirms that Aph treatment essentially delays large (≥300 kb) genes (Figures 3 and S3) ^29^. We therefore focused on those genes for further studies. We used publicly available GRO-seq data from GM06990 lymphoblasts ^43^ to separate large genes in four groups according to their transcription level (Figure 3, GRO-seq) (Methods). We found that genes of the 1^st^ group that comprises the most highly transcribed large genes tend to replicate far earlier than non-transcribed genes of the 4^th^ group. Genes of the other two groups consistently showed intermediate timings (Figure 3, RT).

**Figure 3:**
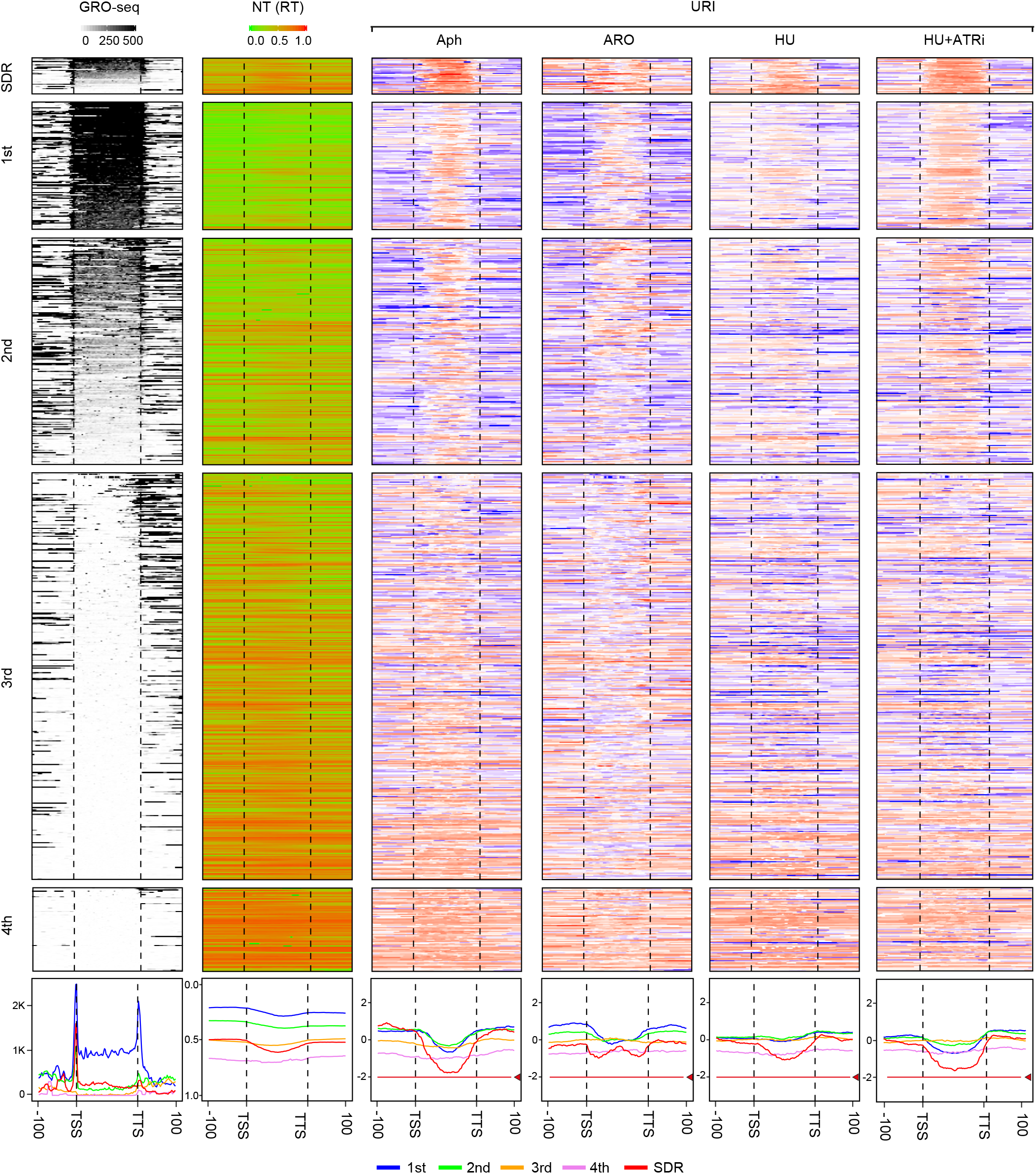
Repli-Seq analyses of URIs along large genes in cells treated with Aph, ARO, HU or HU+ATRi. GRO-seq, replication timing (RT) and URIs along all genes over 300 kb in length and their flanking regions are shown. Cells were treated as in figure 2A. The genes (delineated by dotted vertical black lines) have been rescaled (divided into the same number of bins, independently of their size). Genes hosting SDRs (SDR) are presented above bulk genes. First column: GRO-seq data, the upper bar shows the heatmap color scale. Note that color is saturated for bins mean counts over 500 for visualization purposes. Bulk large genes are divided into 4 groups (1st to 4th) on decreasing transcription levels (Methods). Genes within each group (and within SDR) were also ordered on decreasing transcription levels. Second column: replication timing in non-treated cells (NT (RT)). The heatmap shows the S50 (the moment in S phase when a sequence has been replicated in 50% of the cells) on a scale from 0: earliest to 1: latest. Columns 3 to 6 show URIs in Aph-, ARO-, HU- or HU+ATRi-treated cells (Methods). Heatmap: as in figure 2B. The mean profiles of GRO-seq, RT and URI are presented for each category of genes (lower panels). TSS: transcription start site, TTS: transcription termination site, −100 and +100 correspond to 100 kb flanking regions upstream and downstream of each gene, respectively. Red lines and arrowheads on URI average profiles: as in Figure 2B. Note that the average profile does not reach the −2 threshold for genes containing SDRs because the SDWs are not located at the same place in the different genes.

Analysis of URIs in Aph-treated cells showed strong drops in the body of genes of the 1^st^ and 2^nd^ groups relative to their flanking sequences. URI drops get gradually attenuated in the 3^rd^ and the 4^th^ groups, which can be explained considering that replication initiations are excluded from large genes in proportion to their transcription level ^25^. In addition, we found that late domains hosting some of the genes of the third group and almost all those of the fourth group show URI drops extending over the entire gene and its flanking sequences (Figure 3, Aph). In cells treated with ARO, Aph-induced URI decreases were suppressed, partially in genes of the 1^st^ and the 2^nd^ group and completely in the majority of genes of the 3^rd^ group. The fact that suppression is only partial in the 1^st^ and 2^nd^ groups most probably reflects the temporal order of CyclinA2-CDK1 and CyclinA2-CDK2 assembly during the S phase. Indeed, since CyclinA2-CDK1 complex starts to form around mid-S ^44^, CDK1 inhibition is not expected to impact the early replicating 5′ and 3′ parts of those large genes. Finally, we observed that the replication of genes of the 4^th^ group and their flanking sequences also tend to be advanced in ARO-relative to Aph-treated cells (Paired t-test with the Bonferroni adjustment, p < 2.2e-16) (Figure 3, ARO). Therefore, CDK1 inhibition helps replication of all large expressed genes and, although modestly, of some other non-expressed late replicating domains.

### RO promotes extra replication initiation events in the *FHIT* gene

We used molecular DNA combing (Figures 4A and S4A) (Methods) to decipher the mechanism responsible for URI rescue in cells treated with ARO. We first studied the impact of RO on fork speeds and IODs at the whole-genome level (Figure 4B). We have previously shown that these replication parameters are linearly correlated ^35^. We confirmed this relationship using data obtained with untreated cells and cells treated with various concentrations of Aph, up to 600 nM. In addition, we showed that data from cells treated with RO and ARO align with those obtained with untreated cells and cells treated with Aph 600 nM, respectively (Figure 4B, right panel). RO therefore affects neither fork speeds nor IODs nor the relationships linking these two parameters across the bulk genome.

**Figure 4:**
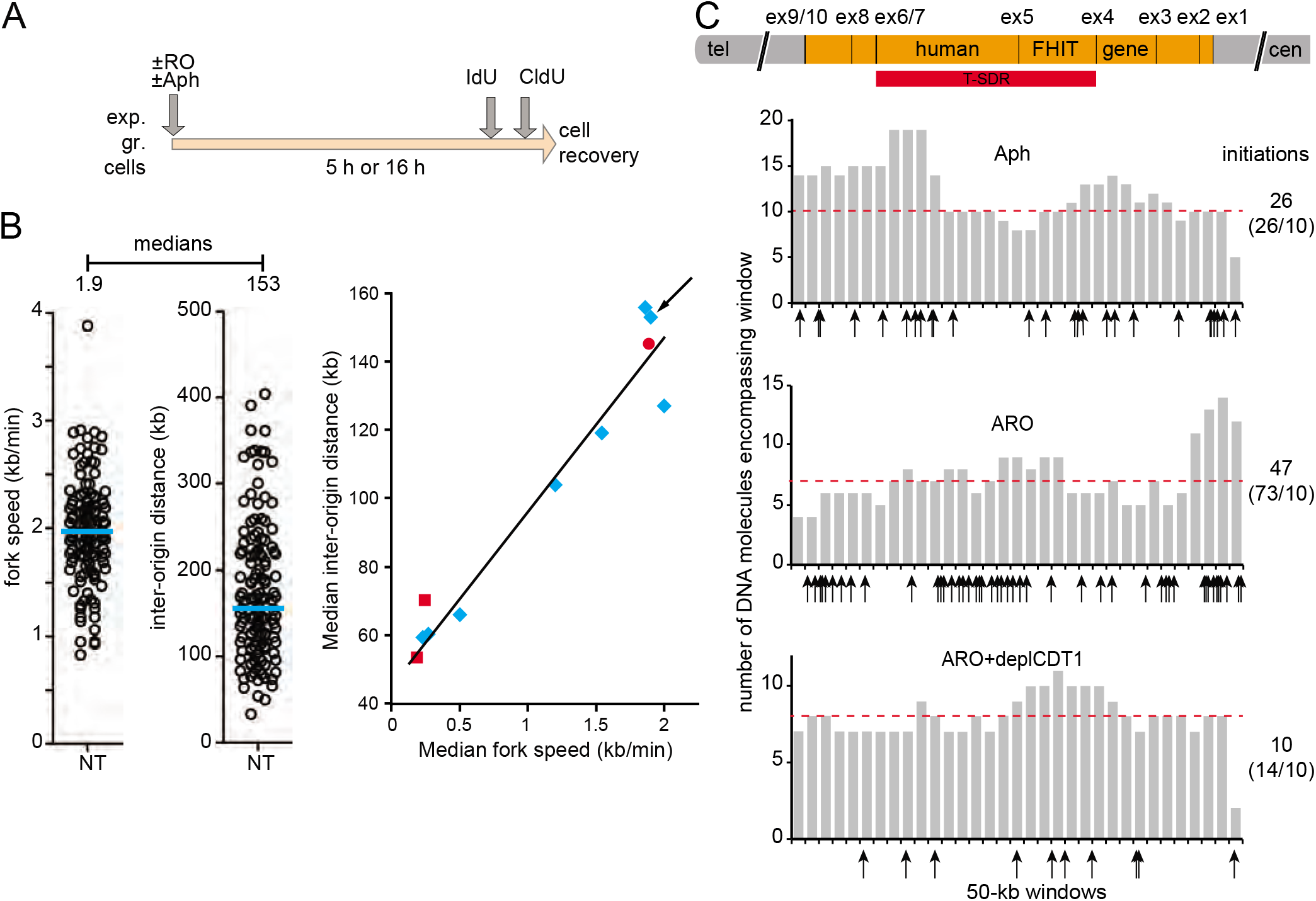
RO does not affect replication dynamics of the bulk genome but promotes extra-replication initiation events in the *FHIT* gene. **A:** Experimental scheme. JEFF cells were grown in the indicated media, labeled with IdU and CldU then treated for DNA combing (Figure S3A) (Methods). **B:** Left panel: Example of fork speed and inter-origin distance analyses in a representative experiment performed with untreated cells. Each open circle corresponds to measurement of one full-size two-color fork (left) or one IOD (right) (Figure S3A). Median fork speed and median IOD were calculated. Right panel: The relationships between median fork speeds and median IODs are presented. The black line was drawn from results of a series of independent biological experiments carried out with untreated cells and cells treated with different concentrations of Aph (blue diamonds). The 3 upper and the 2 lower diamonds correspond to untreated cells and cells treated for 16 h with Aph 600 nM, respectively. The diamond designated by an arrow corresponds to the experiment shown in the left panel. Data obtained for cells treated with ARO (red squares) and RO (red circle) were reported on the graph. Note that the red circle value comes from cells treated with RO for 5 h rather than 16 h because almost no cells remain in S phase after 16 h in the drug (Figures 1B and D). **C:** Mapping of replication initiation events in the *FHIT* gene by DNA combing (Figures S3A and S3B). Cells were treated for 16 h with Aph or ARO as in A or were in addition transfected with siRNAs targeting CDT1 mRNA (deplCDT1) 8 h before ARO treatment. Top diagram: map of the gene on chromosome 3 (3p14.2) with exon (ex) and T-SDR positions. Lower panels: histograms depict coverage, i.e. the number of DNA molecules with replication signal encompassing each 50-kb window along the gene. Median values are indicated by red dotted lines. Arrows indicate the positions of initiation events on the *FHIT* map and their numbers are given on the right with values normalized to coverage of 10 in italics.

We then focused on a region 1.6 Mb-long hosting the *FHIT* gene (Figure S4B). In agreement with the Repli-Seq profiles (Figure 2B), some initiation events were observed along the *FHIT* body after 16 h of treatment with Aph (Figure 4C, Aph). Strikingly, the density of initiation increased approximately by a factor of 3 across the gene in cells treated with ARO (73 events as compared to 27 with coverage normalization to 10) (Figure 4C, ARO). Note that this strong increase cannot be explained by differences in S-phase distribution of cells grown in either media (Figure 1D). Together with the Repli-Seq data, these results show that the URI drop induced in Aph- and suppressed in ARO-treated cells correlates with a burst of extrainitiations taking place in the *FHIT* gene around mid-S phase, and most probably all other large expressed genes. Noticeably, the origins responsible for those extra-initiations are not classical compensation origins, the latter being readily activated in proportion to fork slowing across the bulk genome of cells treated with Aph alone (Figure 4B, right panel).

### Extra initiation events are mandatory to stabilize CFSs

We then asked whether extra initiation events observed in ARO are causal to suppression of Aph-induced instability of FRA3B in JEFF cells. In eukaryotic cells, origin setting and firing requires the activity of CDC7-DBF4 ^45^. We therefore treated the cells with PHA-767491 (PHA), a potent CDC7 inhibitor with more than 20-fold selectivity against CDC7 than against CDK1 or CDK2 ^46, 47^. To confirm this selectivity, we checked the status of MCM2 at residues specifically phosphorylated by CDKs or CDC7. We choose to use PHA at 6 μM, a concentration that strongly reduces CDC7-dependent phosphorylation of S40 and S53 without impacting CDK-dependent phosphorylation of S27 and S41 (Figure S4A).

We first determined how treatments with Aph, RO, PHA 6 μM and their combinations (Figure 5A) impact cell cycle progression (Figure 5B). We observed that cells grown in the presence of PHA accumulate in G1, showing that once JEFF cells are engaged in the S phase, CDC7-DBF4 activity is required neither for replication completion nor for cell cycle progression up to the following G1 phase. In RO+PHA the cells accumulated in G1 because of PHA and in G2 because of RO. We however observed that cell progression across late S phase is far slower in RO+PHA than in RO or PHA alone (Figures 5B and 1D). Cell accumulation upon growth in Aph+PHA or in ARO+PHA was consistent with these conclusions taking into account that Aph strongly slows cell progression all along the S phase (Figure 5B). We then studied metaphase chromosome stability as above (Figure 5C). Determination of total breaks in untreated cells and cells treated with Aph or ARO confirmed the results shown in figure 1E. In cells treated with PHA alone or with RO+PHA, break frequencies were comparable to those found in untreated cells. Strikingly, break frequencies in cells treated with Aph, Aph+PHA or ARO+PHA were similar, showing that CDC7-DBF4 activity is required during the S phase to allow suppression of CFS instability by RO. These results confirm that, in contrast to classical compensation origins, extra-origins involved in completion of large gene replication in cells treated with ARO are not ready to start, raising the question whether they are built *de novo* during the S phase, possibly from new pre-RC.

**Figure 5:**
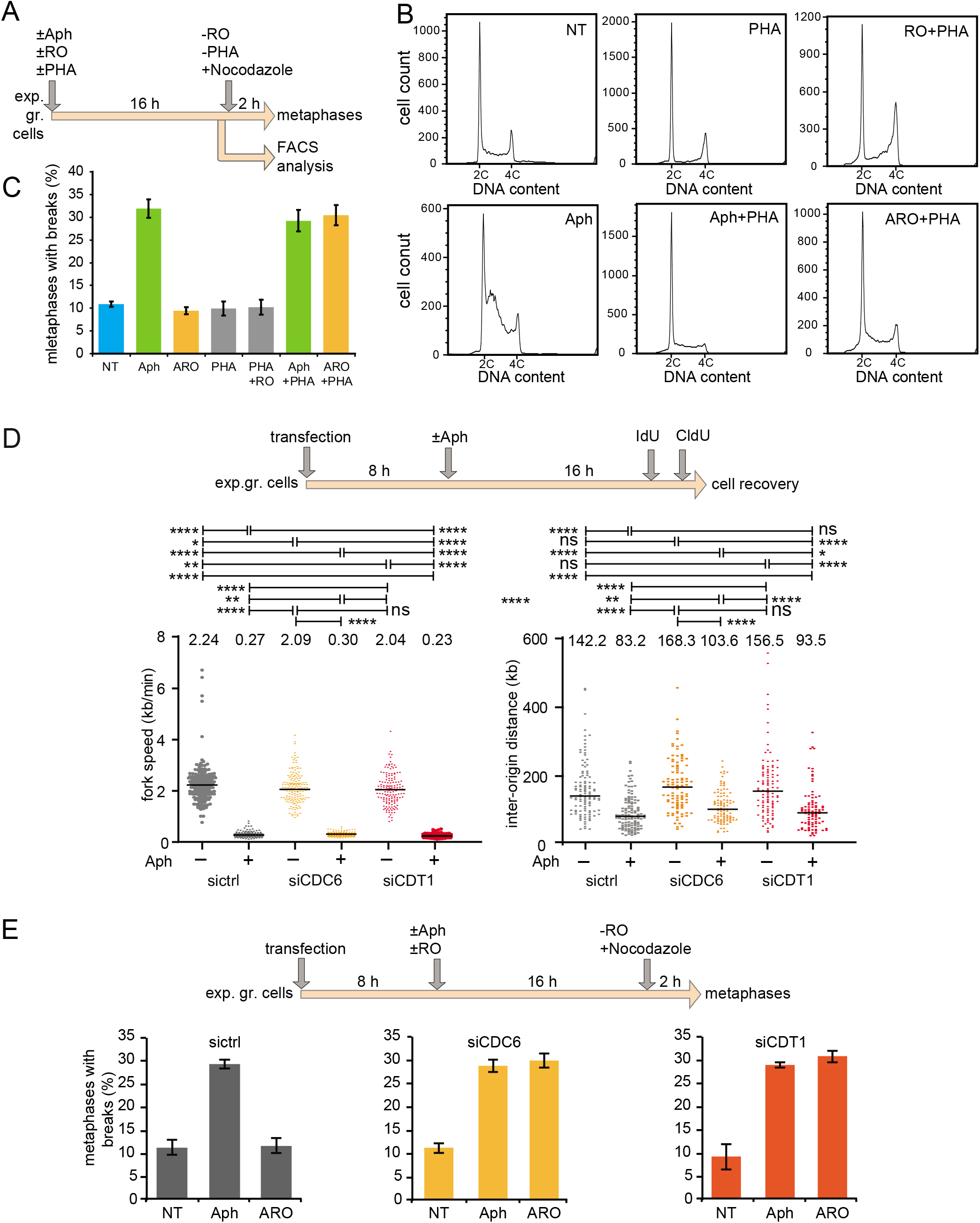
Stability of CFSs in ARO treated cells relies on extra initiation events, CDC6 and CDT1. **A-C:** Exponentially growing (exp. gr.) JEFF cells were incubated in the presence or absence of Aph and/or RO and/or PHA (6 μM) (A) and prepared for FACS analysis (B) or determination of total break frequency as in figure 1 (C). **D:** Experimental scheme is shown. Cells were transfected with siRNAs targeting CDC6 or CDT1 mRNAs (siCDC6, siCDT1) or with a control siRNA with no known target (sictrl). Fork speed and IOD were determined for cells treated as indicated. Results are presented as in figure 4B. Significance is shown at the top (horizontal lines): ns not significant, * p<0.05, ** p<0.01, **** p<0.0001. E: Cells were transfected with siCDC6, siCDT1 or sictrl RNA as in D, and then treated as indicated. Total breaks were determined as in figure 1E. Experiments shown in C and E were carried out twice and error bars represent standard deviation.

### Stability of CFSs in cells grown in ARO relies on CDC6 and CDT1

Redundant mechanisms normally inhibit building of pre-RCs during the S and G2 phases. Prominent among these mechanisms are CDK1-dependent degradation of CDT1 and nuclear exclusion of CDC6 ^45^. To determine if these pre-RC components play a role in CFS rescue in cells grown in ARO, JEFF cells were transfected with siRNAs targeting CDC6 or CDT1 mRNAs (siCDC6, siCDT1) or with a control siRNA with no known target (sictrl). Western blot analyses showed that both proteins are severely depleted 8 h post-transfection and that depletion persists at least up to 24h (Figure S5B). We also studied the impact of transfection *per se* on cell cycle progression (Figures S5C-S5H). Whatever the siRNA used, including the sictrl, we found that JEFF cells transfected in G1 phase enter S phase extremely slowly. Indeed, only a small fraction of G1 cells has progressed to S phase 24 h post-transfection. In contrast, cells transfected in S or G2 phase displayed a lag of only 6 h (Figures S5C-S5H). Using the time frame described in figure 5D, this unexplained phenomenon permitted us to study cells that have been depleted of pre-RC factors during the S phase. Molecular combing analysis of the replication dynamics across the bulk genome of cells transfected with the sictrl, siCDC6 or siCDT1 showed that fork speed and IODs are marginally, if at all, impacted by depletion of either pre-RC factor (Figure 5D). Since availability of CDC6 and CDT1 is mandatory in G1 phase to sustain efficient pre-RC building, these results confirm that cells were engaged in the S phase prior to depletion.

The frequencies of total breaks were then determined in cells transfected with the sictrl, siCDT1 or siCDC6 and further grown either in normal medium, Aph, RO or ARO (Figure 5E). Cells transfected with the sictrl displayed a break frequency similar to that found in non-transfected cells grown in the same media (Figure 1E), which shows that transfection *per se* does not affect chromosome integrity under the experimental conditions used here. Strikingly, CDC6 or CDT1 depletion completely prevented RO-dependent rescue of Aph-induced breaks (Figure 5E). These results therefore strongly support the idea that pre-RC can be reloaded during the S phase in the absence of CDK1 activity, rescuing CFS stability upon fork slowing.

### CDT1 is required to reset the initiation program across the *FHIT* gene in ARO-treated cells

Focusing on CDT1, we used molecular combing to study the density of initiation events along the *FHIT* gene. Strikingly, comparison of ARO-treated cells depleted or not of CDT1 revealed a drastic decreased in the number of initiation events along the gene in depleted cells (Figure 4C, lower panel). For median coverage normalized to 10, 14 initiation events were observed in CDT1 depleted cells as compared to 73 in proficient cells. With similar normalization to 10 times coverage, we have previously observed 12 initiation events in untreated JEFF cells and 31 upon treatment with Aph 600 nM (26 here) ^48^. Comparison of the number of initiation events in ARO-treated cells depleted of CDT1 to that found in untreated cells (14 and 12, respectively) suggests that CDT1 depletion completely prevents stress-induced extra initiation events along the *FHIT* gene, *i.e*. those due to RO and those due to Aph. Therefore, although Aph elicits weak DRC activation, it may promote firing of a few extra-origins in the *FHIT* body, but not enough to rescue FRA3B stability. We concluded that building of functional extra-origins occurs during the S phase along *FHIT,* and most probably all other large expressed genes, in proportion to the degree of CDK1 inhibition.

### Stringent checkpoint activation by HU protects CFS integrity

We reasoned that if the stringency of CDK1 inactivation dictates instability of CFS, they should be poorly, or not at all, destabilized by replication stresses that trigger a robust DRC response. To check this hypothesis, we treated JEFF cells for 16 h with HU 150 μM. This concentration of HU induced g-H2AX formation during the S phase (Figure 6A), and strong CHK1 and p53 phosphorylation (Figure 6B). However, HU 150 μM slowed fork progression to the same extent as Aph 600 nM (Figures 6C and 4B). Cytogenetic analyses showed that HU treatment results in chromosome breaks (Figure 6D). Total breaks were induced approximately 1.6 folds more efficiently by HU-than by Aph treatment. In striking contrast, the frequency of breaks at FRA3B was approximately 3 folds lower in HU than in Aph and FRA16D was not at all destabilized in HU, which strongly supports our hypothesis.

**Figure 6:**
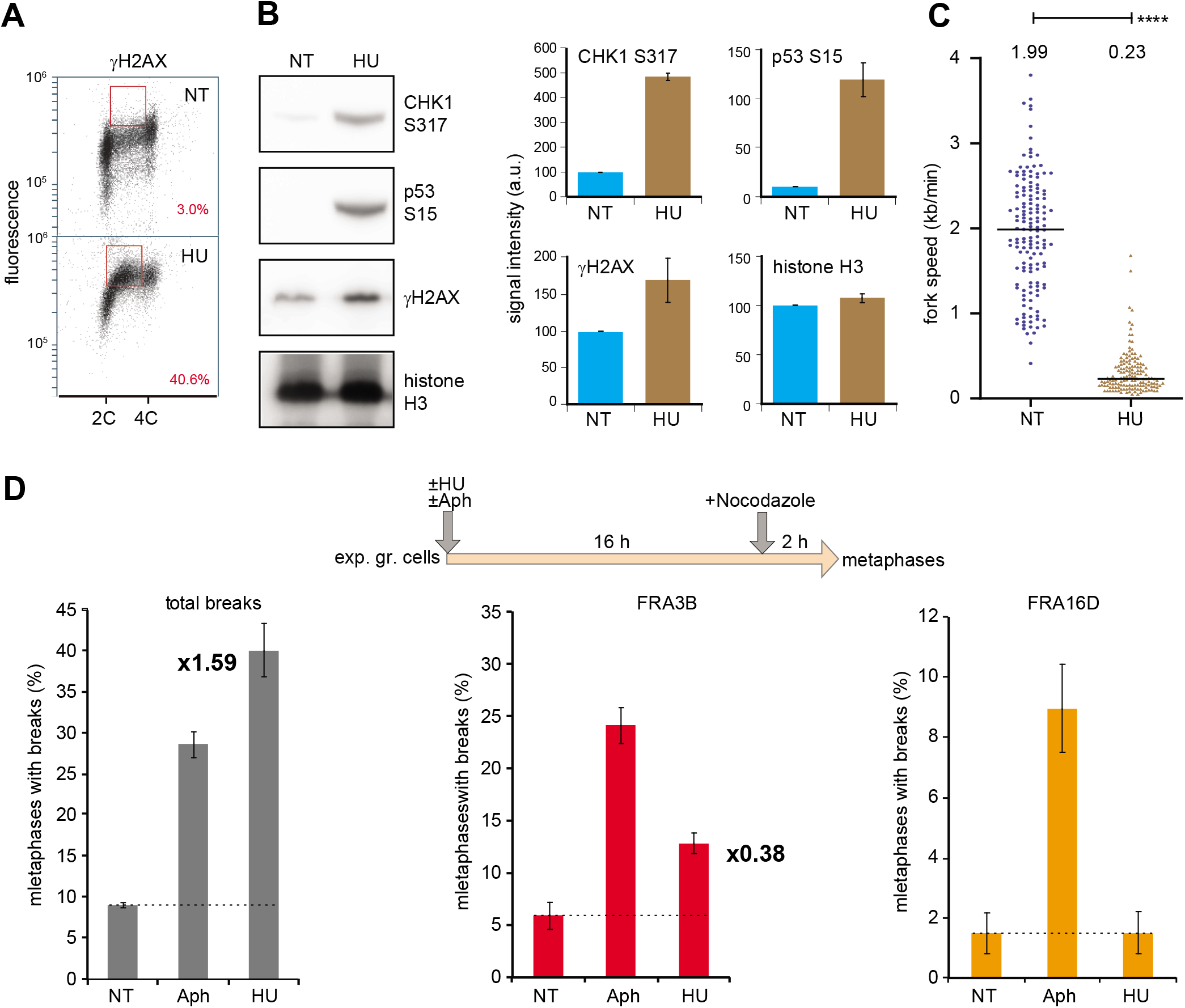
Checkpoint activation by HU protects CFS integrity. **A:** JEFF cells untreated (NT) or treated with HU 150 μM (HU) for 16 h were analyzed by FACS with antibodies anti-histone γH2AX. Percentages of labeled S-phase cells (present in the red squares) are indicated in red. **B:** Cells were treated as in A, total protein extracts were prepared and analyzed by Western blotting with antibodies anti-phospho-ser317 CHK1, anti-phospho-ser15 p53, anti-histone γH2AX or anti-histone H3 (as a loading control). A representative experiment is shown (left panel). Signals on digital images were quantified with Image Gauge (right panels). **C:** Cells were treated as in A, and fork speeds were determined and presented as in the left panel of figure 3B. **D:** Cells were treated as indicated. Experimental procedures are described in figure 1E. Comparison of break frequency between Aph and HU: fold increase or decrease was calculated after background subtraction (dotted lines) and is indicated for total breaks and FRA3B (cannot be calculated for FRA16D). Experiments shown in B and D were carried out twice and error bars represent standard deviation.1^st^ and 2^nd^ groups differs in cells treated with HU or ARO, essentially because drop attenuation starts earlier in the S phase of HU-treated cells. This observation suggests that, in addition to CDK1, DRC effector(s) functioning before mid-S contribute to stress-induced advancement of large gene replication in HU-treated cells. In order to definitely establish causal relationship between modest URI drop in HU and checkpoint activation, we analyzed cells co-treated with HU and ATRi. We found that ATR inhibition does potentiate the drop, especially for SDRs and the 1st and 2nd groups (Figures 2B-C and S2F, and Figure 3).

We then performed Repli-Seq analysis of cells treated for 16 h with HU 150 μM. We observed a burst of DNA synthesis across the genes *FHIT* and *WWOX* that strongly limits URI drop in the former and completely prevents it in the latter (Figure 2B). These results therefore perfectly agree with the impact of HU treatment on FRA3B and FRA16D stability (Figure 6D). Global analysis of SDRs and T-SDRs confirmed genome-wide that URI decreases are strongly attenuated in HU-relative to Aph-treated cells (Figures 2C, 3 and S2F). Noticeably, the SDRs and T-SDRs of HU-treated cells displayed URI profiles comparable to that of the bulk of non-fragile late replicating DNA (Figure 2C). Finally, we studied the effect of HU treatment on the replication of genes longer than 300 kb (Figure 3). Data showed weak, if at all, URI drop in genes of all four groups. In addition, we observed that the kinetics of URI drop in genes of the 1^st^ and 2^nd^ groups differs in cells treated with HU or ARO, essentially because drop attenuation starts earlier in the S phase of HU-treated cells. This observation suggests that, in addition to CDK1, DRC effector(s) functioning before mid-S contribute to stress-induced advancement of large gene replication in HU-treated cells. In order to definitely establish causal relationship between modest URI drop in HU and checkpoint activation, we analyzed cells co-treated with HU and ATRi. We found that ATR inhibition does potentiate the drop, especially for SDRs and the 1st and 2nd groups (Figures 2B-C and S2F, and Figure 3).

## Discussion

Here we show that stringent CDK1 inhibition suppresses stress-induced CFS instability. To understand how, we compared the replication dynamics of JEFF cells treated with Aph or ARO to that of cells in which CDK1 is, in addition, inhibited by RO. We found that Aph triggers under-replication of large origin-poor genes hosting CFSs, in complete agreement with previously reported results ^29^. In ARO-treated cells, Repli-Seq revealed a burst of newly synthesized DNA implemented from mid S phase in the body large of fragile genes. This synthesis permits completion of CFS replication during the S phase, which well explains their rescue. In striking contrast, it was recently reported that replication completion of CFSs occurs during mitosis in cancer-derived U2OS, HeLa and HCT116 cells grown in the very same conditions, *i.e*. for 16-24 h in ARO ^42, 49, 50^. This phenomenon, called mitotic DNA synthesis (MIDAS), was however considerably attenuated or totally absent in cells closer to normal. We propose that, in primary fibroblasts such as HS68 ^49^ and MRC5 (this work) as well as in immortalized HBE bronchial epithelial cells ^50^, RPE1 retina epithelial cells ^42^ and lymphoblasts (this work), CDK1 inhibition stimulates replication completion of CFSs during the S phase while some cell lines derived from advanced tumors are deficient in this DRC response. Altogether, results suggest that MIDAS is used as a backup mechanism in cells that fail to rescue CFS replication during the S phase, and that such failure strongly contributes to generate the genomic alterations found in cells of various cancers.

We also treated JEFF lymphoblasts with HU 150 μM, a drug concentration leading to the very same level of fork slowing than Aph 600 nM, the concentration used in experiments reported here. In contrast to Aph, HU triggered stringent DRC activation in JEFF cells. This response most probably results from damages elicited by the severe imbalance of the dATP/dTTP pools induced by the drug ^35^ rather than from fork slowing *per se*. We therefore asked whether this concentration of HU destabilizes CFSs. In agreement with previous reports ^51, 52^, we observed breaks in metaphase chromosomes, part of them at CFSs. However, we found that only a subset of the Aph-sensitive CFSs displays breaks in HU, and at far lower frequency than in Aph. These latter results are consistent with Repli-Seq data showing that a burst of DNA synthesis occurs in mid S phase along the body of fragile genes in HU-treated cells, which alleviates or completely prevents their under-replication. In addition, we found that this burst is strongly attenuated in cells co-treated with HU and VE822 to inhibit ATR activity. Together, results therefore show that DRC-induced CDK1 inhibition advances replication completion of large genes depleted of origins by transcription.

To decipher the mechanism responsible for stimulation of CFS replication upon CDK1 inhibition, we studied the impact of RO on replication speed and initiation density across the whole genome and along the *FHIT* gene using molecular combing. Comparison of cells treated with Aph or with ARO showed that Aph dictates fork speed and IODs in the bulk genome, independently of RO. In striking contrast, we observed a strong increase in the density of initiation events across the gene in cells treated with ARO. The corresponding extra-origins therefore behave differently from classical compensation origins, the latter being readily activated in the bulk genome of cells treated with Aph alone. Whatever their nature, firing of extra-origins well explains the burst of newly synthesized DNA across *FHIT* in cells grown in ARO. We then asked whether this local effect of RO on origin density correlates with a reduced expression of large genes hosting CFSs. We showed that Aph-, RO- or ARO-treatments do not substantially impact the transcription level of the 3 fragile genes we studied. These results therefore support our previous conclusion that large gene instability primarily relies on transcription-induced origin paucity rather than on conflicts between the transcription and replication machines ^29, 48^. Remarkably, the kinetics of replication completion along CFSs in cancer cells undergoing MIDAS also offers strong support to this conclusion ^49^.

In order to determine if the effect of RO is limited to genes hosting CFSs we extended our analysis to all genes displaying replication delay upon Aph treatment. These genes share the following properties: (i) they are continuously transcribed over at least 300 kb but may be expressed from very low to moderate levels, as expected for genes of that size ^29^; (ii) they terminate replication in the first half of S phase in untreated cells and in the second half upon Aph treatment, explaining why they display neither under-replication nor instability; (iii) their delayed replication is only partially rescued in ARO, which can be explained considering that they start to replicate far before mid S, a time at which cyclin A-CDK1 is inactive; (iv) their replication is weakly, if at all, delayed in HU-treated cells. Therefore, rescue of replication delay upon DRC activation is not limited to genes hosting CFSs. Moreover, these results confirm that late replication completion is mandatory to set CFSs in addition to long initiation-poor regions ^28^.

Hypothesizing that transcription may either displace the pre-RCs from their sites of loading or favor their degradation, we asked whether CDK1 inhibition allows unscheduled building or repositioning of pre-RCs during the S phase. We reasoned that, in contrast to repositioning, building requires that proteins involved in pre-RC loading remain available and functional in S phase. Various observations support this possibility, notably: (i) CDK1 phosphorylates CDC6, which promotes its export out of the nucleus ^45^; (ii) multiple pathways limit CDT1 availability during the S-phase, including degradation mediated by two E3 ubiquitin ligases, CRL4^Cdt2^ and CRL1^Skp2^, the latter pathway relying on CDT1 phosphorylation by S phase CDKs ^45, 53^; (iii) several pre-RC components, notably CDC6 and CDT1, contain intrinsically disordered regions that promote formation of condensates, helping MCM recruitment to the chromatin. Strikingly, a recent work showed that the S-CDKs disrupt these condensates ^54^. Here we showed that depletion of CDC6 or CDT1 during the S phase prevents CFS rescue in ARO-treated cells. Focusing on CDT1, molecular combing experiments consistently showed that extra-initiation events taking place along the *FHIT* gene body no longer occur in ARO-treated cells depleted of the protein in S phase. We also showed that inhibition of CDC7-DBF4 activity during the S phase also prevents RO-dependent rescue of CFS stability. We conclude that firing of new origins, build on pre-RCs loaded during the S phase upon CDK1 inhibition, relieves transcription-induced origin paucity of genes hosting CFSs, and probably of all other large expressed genes.

To maintain genome stability, the replication process should duplicate the genome once and only once. To avoid re-replication, the level of pre-RC components is tightly regulated, particularly that of CDT1 ^45^. Surprisingly, although this protein remains functional during the S phase in the absence of CDK1 activity, genome stability was not altered in human lymphoblasts and rodent fibroblasts studied here. In agreement with a previous report ^39^, we indeed found that cells accumulated in G2 phase upon RO treatment as well as cells released in normal medium that undergo mitosis then progress across the next cell cycle display neither breaks in mitotic chromosome nor g-H2AX accumulation in interphase. A possible explanation comes from reports showing that while CDT1 degradation *via* CRL4^Cdt2^ ubiquitin ligase is mandatory to prevent re-replication ^55, 56^, its CDK-dependent degradation *via* CRL1^Skp2^ is not ^57, 58^. In line with the protective effect expected from DRC activation, we propose that CDT1 accumulation resulting from inactivation of the CDK1-dependent CRL1^Skp2^ pathway is sufficient to permit targeted pre-RC building during the S phase but not to sustain re-replication.

It is generally admitted that the main role of the DRC is to prevent cells with incompletely replicated and/or damaged DNA to enter mitosis. However, our finding that DRC activation fosters replication termination of late replicating large expressed genes, and some other late replicating origin-poor domains, reveals a yet unsuspected role of DRC activation in the protection of genome stability. Advancement of the replication time of sequences at risk of under-replication indeed provides decisive support/alternative to DRC-dependent delay to mitotic onset. However, CFSs remain under-replicated in a particular window of stress in which the level of CDK1 inhibition is insufficient to induce enough extra-initiations to rescue CFSs and to prevent mitotic onset. Noticeably, these low levels of stress elicit only a few breaks in a limited fraction of the cells. We therefore propose that this window of stress plays a critical role in the balance between protection of genetic information and generation of beneficial somatic diversity in some cell lineages. This phenomenon is best exemplified in neuronal stem-progenitor cells in which many large genes hosting CFSs are expressed, supporting the idea that recurrent breaks taking place in these genes drive neuronal diversity and normal neurological development ^38^.

## Methods

### Cell culture

JEFF cells (human B lymphocytes immortalized with the Epstein–Barr virus) were grown as previously described ^48^. Culture medium was supplemented with 3 μM of each 2′-deoxyadenosine, 2′-deoxycytidine, 2′-deoxyguanosine and thymidine (Sigma) for experiments with PHA767491 (Selleckchem, S2742). Aphidicolin (A0781), RO (CDK1 Inhibitor IV - Calbiochem; 217699), VE-821 (Sigma-Aldrich SML1415) and hydroxyurea (Sigma-Aldrich H8627) were obtained from Merck. MRC5 cells (primary human lung fibroblasts) were grown at 37 °C in a humid atmosphere containing 5% CO_2_ and 3% O_2_ in MEM medium containing 10% fetal calf serum, non-essential amino acids, 1 mM sodium pyruvate, 2 mM glutamine and 1% penicillin and streptomycin.

### Fluorescence-activated cell sorting (FACS)

See supplemental Table I for a detailed description of the primary and secondary antibodies used in these studies. Analyses of BrdU pulse-labeled JEFF cells were carried out using rat anti-BrdU and chicken anti-rat Alexa 488 antibodies as described ^35^. For MPM2-positive cells, cells were fixed in 75% ethanol overnight at 4°C, washed with phosphate buffered saline solution (PBS) and incubated for 45 min at room temperature with anti-MPM2 antibodies in PBS containing 1% bovine serum albumin and 0.5% Tween 20. After wash with PBS, they were incubated for 30 min at room temperature with goat anti-mouse immunoglobulins-Alexa 488 antibodies in the same buffer. After PBS wash, cells were resuspended in PBS containing RNase A (25μg/mL) and propidium iodide (50 μg/mL). Samples were analyzed using a BD Biosciences LSRII flow cytometer and data were analyzed using FlowJo v8.7.3 software. For FACS analysis of gamma H2AX −positive cells, cells were fixed and processed as above using mouse monoclonal anti-phospho-histone H2AX and goat anti-mouse immunoglobulins-Alexa 488 antibodies. For Repli-Seq experiments cells were sorted into 6 fractions (G1/S1, S2, S3, S4, S5, S6/G2M) as illustrated in figure S2A.

### Cytogenetics

Analysis of total breaks on Giemsa-stained chromosomes was done as described ^29^. Preparation of BACs selected from the human genome project RP11 library and labeling of probes was carried out as previously described ^61^. Fluorescence in situ hybridization (FISH) on metaphase chromosomes and immunofluorescence revelation of FRA3B-, FRA16D-, FRA1L-, or FRA3L-specific probes were carried out as previously described ^25, 48^. BACs used were: FRA3B: RP11-32J15, RP11-641C17, RP11-147N17; FRA16D: RP11-105F24, RP11-57106; FRA1L: RP11-88425, RP11-729G19; FRA3L: RP11-59m6, RP11-10915, RP11-739D3. In order to reveal the centromeres of the chromosomes of interest, probes corresponding to the alpha satellite sequences of chromosome 3 (Aquarius probes LPE 03 R) or of chromosome 16 (Aquarius probes LPE 16 R) (Cytocell) were used.

### Immunofluorescence

GMA32 cells were grown in 2 cm-Petri dishes. After treatments, they were washed with PBS, fixed with 4% formaldehyde for 5 min at room temperature, washed 3 times in PBS, permeabilized by incubation for 5 min at room temperature in PBS containing 0,5% Triton X-100 (PBS-T) and washed again with PBS. They were then incubated for 30 min at room temperature in PBS containing 1% bovine serum albumin (BSA), then for 1 h at room temperature with rabbit polyclonal anti-gamma H2AX antibodies in the same buffer. After washing twice with PBS-T and once with PBS, they were incubated for 30 min at room temperature with goat anti-rabbit immunoglobulins-Alexa 594 antibodies in PBS/BSA. After washes with PBS-T and PBS as above, they were mounted in Vectashield DAPI (Vector Laboratories, H-1200) under coverslips. After treatments, JEFF cells were collected and resuspended in PBS at 400 000 cells/mL. Two hundred μL were loaded in chambers of a Cytospin centrifuge and cells were deposited onto vertical microscope slides by centrifugation for 3 min at 700 rpm. Cells were fixed with 2% formaldehyde in PBS, washed with PBS and permeabilized as described for GMA32 cells. Treatments with primary (mouse monoclonal anti-phospho-histone H2AX) and secondary (goat polyclonal anti-mouse immunoglobulins-Alexa 594) were carried out and slides were mounted as described above.

### DNA combing

Labeling of neo-synthesized DNA with iododeoxyuridine and chlorodeoxyuridine, combing of DNA on silanized coverslips, design of the *FHIT* Morse code, FISH on combed DNA and immunofluorescence detection, determination of fork speed, initiation density, inter-origin (IOD) distances and coverage and statistical analysis of molecular combing data have been previously described ^48, 60^

### SiRNA transfection

Transfections were carried out with the Nucleofector device (LONZA), according to the manufacturer’s instructions as reported ^35^. After electroporation, the cells were immediately diluted in culture medium at 37 °C. SiRNAs targeting human mRNAs were purchased from Invitrogen. SiCDT1: Stealth siRNAs (Set of 3) HSS129994, HSS188618, HSS188619; siCDC6: Stealth siRNAs (Set of 3) HSS101647, HSS101648, HSS101649. For control experiments, cells were transfected with the AllStars Negative Control siRNA (SI03650318, Qiagen) referred as the sictrl throughout the article.

### RNA isolation and quantification

See supplemental Table II for the sequences of the primers used in these studies. Preparation and quantification of nascent RNA have been reported ^29^. For quantification of primary transcripts, total RNA was extracted using RNeasy Mini Kit (Qiagen) according to the manufacturer’s instructions. Chloroform extracted RNA was treated with RNase free recombinant DNase I (047 716 728 001, Roche) for 1 h at 37°C using 2 units/μg of RNA. The reaction was stopped by adding 10 mM EDTA and RNA was purified by extractions with phenol/chloroform (1/1) and then chloroform. It was ethanol precipitated, washed with ethanol 70%, redissolved in RNase-free water and quantified by spectrophotometry. RNA (1 μg) was reverse transcribed using the SuperScript III First-Strand Synthesis Kit (Invitrogen 18080). Primary transcripts were quantified by real-time quantitative PCR (qPCR) using intronic primers. Reactions were carried out using the 7500 Real time PCR system (Applied Biosytem) with the SYBR green PCR Master Mix (Applied Biosystem). Each reaction was performed in duplicate. The absence of genomic DNA contamination was assessed using RNA samples incubated in reverse transcriptase-free reactions. Results were normalized using cyclophilin B (PPIB) RNA as endogenous control and are presented relative to the level in untreated cells (NT).

### Protein extraction and Western blotting

Total protein extracts were obtained from cell pellets directly resuspended in the loading buffer (Biolabs B7709S) containing 1% SDS, 10 mM DTT and 10 mM MgCl_2_ (25 μL/10^6^ cells) and then incubated for 30 min at room temperature in the presence of 75 U of benzonase (Millipore D0017) in order to digest DNA. Before electrophoresis, the samples were heated for 5 min at 95 °C. Proteins were analyzed by electrophoresis in pre-cast gels 4-12% polyacrylamide-SDS gels (Life Technologies NP0322) using the corresponding migration buffer (MOPS NP0001). Transfer onto nitrocellulose membranes was carried out using program 3 of an Iblot machine (Life Technologies) and the associated transfer kit (Life Technologies IB301001). The membranes were then saturated by incubation for 1 h at room temperature in PBS containing 0.05% Tween 20 and 5% BSA. Incubations with the various primary antibodies and with secondary antibodies coupled to horseradish peroxidase (see supplemental Table I) were carried out overnight in the same buffer at 4 °C and 1 hour at room temperature, respectively. After incubation with antibodies, membranes were washed 3 times for 10 min in PBS containing 0.05% Tween 20. The chemiluminescence signals were revealed using the WesternBright ECL kit (Advansta K-12045). Signals on digital images were quantified with the Image Gauge software (Fujifilm).

### Chromatin immunoprecipitation

Chromatin preparation: JEFF cells were resuspended in PBS at 10^7^ cells per mL and fixed by incubation for 10 min at room temperature with stirring in the presence of 1% formaldehyde. Formaldehyde was neutralized by adding 1.25 M of glycine and the cells were washed twice with cold PBS. They were resuspended at 10^8^ cells per mL in lysis buffer (50 mM Hepes pH6.8, 150 mM NaCl, 1 mM EDTA, 1% Triton X-100, 0.1% deoxycholate of sodium and 0.5% SDS) containing protease inhibitors. After 30 min at 4 °C., the lysate was diluted 10 times with lysis buffer and subjected to sonication for 15 min in a refrigerated water bath (Bioruptor, Diagenode). Part of the sonicated chromatin was treated with proteinase K and RNase A and analyzed by electrophoresis in a 1.5% agarose gel to control the size of the DNA fragments (200-500 bp). Another part of the prepared chromatin did not undergo immunoprecipitation, serving as reference to determine the abundance of sequences of interest in the starting chromatin (input). Immunoprecipitation: 50 μL of Protein A beads suspension (Millipore 16-157) were resuspended in PBS containing 2% BSA and 5 μg of non-immune rabbit polyclonal immunoglobulins (ab37415-5) or 3 μg of antibodies directed against the phosphorylated form of serine 5 of the C-terminal domain of RNA polymerase II (ab5131), for control tubes and test tubes respectively. After incubation for 2 h at 4 °C, chromatin corresponding to 2×10^6^ cells was added and incubation was pursued overnight at 4 °C with stirring. Beads then underwent a series of successive washes at 4 ° C (i) in the lysis buffer; (ii) in the lysis buffer containing 500 mM NaCl; (iii) in LiCl buffer (0.25 M LiCl, 0.5% NP40, 1 mM EDTA, 10 mM Tris-Cl pH8.0, 0.5% sodium deoxycholate); (iv) in TE buffer (10 mM Tris-Cl pH8.0, 1 mM EDTA). Reversion of coupling to formaldehyde and quantification: 125 μL of reversion buffer (25 mM Tris-Cl pH8.0, 5 mM EDTA, 0.5% SDS) were added to the tubes which were incubated for 20 min at 65 °C with shaking. After centrifugation to remove the beads, the supernatants were supplemented with 100 μg/mL of proteinase K and incubated for 2 h at 65 °C. RNase A (15 μg/mL) was then added and incubation was continued for 30 min at 37 °C. DNAs were purified using the Quiaquick PCR purification Kit (Qiagen 28104) and eluted in 100 μL of water. The sequences of interest were quantified by qPCR using intronic primers (supplemental Table II). Amounts of precipitated DNA were determined using a reference curve obtained with increasing known quantities of sonicated DNA extracted from JEFF cells. Obtained values were normalized to the DNA quantities determined in the corresponding input samples. Final results were expressed as fold enrichments, calculated as the ratio of the standardized amount obtained for the test to that obtained for the corresponding negative control.

### Repli-Seq data processing

The raw Repli-Seq data, their processing and URI calculation were performed as described in^29^ with modifications as follows. Technical biases due to PCR amplification were removed and then further reduced by subtracting to each phase a control BrdU IP performed on cycling unsorted cells. The center of the first peak of the read number distribution was then identified as the noise threshold, so that the center of the first peak of each phase was centered at zero and negative values were removed. A second normalization step is based on a reference FACS profile reporting the relative positions of the gates used for the FACS sorting (Figure S2A) and a simulation of the S phase progression based on ^59^ (Figure S2B). Thanks to this simulation we were able to calculate the amounts of newly replicated DNA (based on the total number of active forks) in function of the percentage of replicated DNA in each fraction (Figure S2C). The 6 Repli-Seq fractions were therefore re-sized to the total number of 90 M reads and distributed between each phase proportionally to the mean number of active forks replicating the genome within each fraction.

### Genome wide data analysis

GRO-seq (GEO: GSM1480326) signal intensity was computed with the total number of counts from both strands over 10 kb bins. For genes larger than 100 kb, a weighted median intensity has then been calculated for each gene, and used to divide them into 4 groups based on GRO-Seq level. The 1^st^, 2^nd^ and 3^rd^ group contain transcribe genes with respective weighted median GRO-seq signal of [1 - 0.66), [0.66 - 0.33) and [0.33 – 0] percentiles, respectively. The percentiles have been calculated after excluding genes with weighted median GRO-seq signal equal to zero that we consider as non-transcribed genes and belong to the 4th group. The matrixes for the heatmaps were produced using Enriched Heatmap R (package v1.12.0, R v3.5.2) where the mean intensity (w0) of GRO-seq, URI and RT signals was resizing the gene bodies over a fixed region and including 100 kb upstream and downstream of each gene. Profiles report the mean signal in each bin.

## Acknowledgments

The three teams contributing to the work (M.D., C-L.C., C.T.) have been supported by the Fondation pour la Recherche Médicale (FRM) (programme DBI20131228560). M. D.’s team is also supported by the Institut National du Cancer (INCa) (subvention AAP INCa 2017 −PLBIO17-194). The M.D.′ and C-L.C.’s teams are supported by the Agence Nationale pour la Recherche (ANR) AAPG 2019 “TELOCHROM”. The C-L. C.’s team is in addition supported by the ANR "ReDeFINe", the INCa PLBIO19 076 and by grants from the Curie Institute YPI program, the ATIP-Avenir program from CNRS and Plan Cancer from INSERM, CNRS 80|Prime inter-disciplinary program. S.G. would like to thank ATIP-Avenir and Plan Cancer for his post-doc fellowship. S.E.-H. was supported by the FRM program. D.A. was supported by a Ph-D fellowship from the Ligue Nationale Contre le Cancer.

We thank Aline Renoult for help with some gamma-H2AX foci experiments. We acknowledge the Imaging and Cytometry Platform (UMS 3655 CNRS/US 23 INSERM) of Gustave Roussy Cancer Campus for assistance with cell sorting, and Michael Schertzer for critical reading of the manuscript.

## Author contribution

M.D. and O.B. conceived the project and wrote the paper. C.-L.C., S.G. and C.T. provided critical revision of the manuscript and contributed to figure preparation. O.B. contributed to and directed D.A., M.S. and A.-M.L. bench-work, and analyzed the results. D.A., S.K., M.S., and A.-M.L. contributed to biological experiments, including cell culture, cell sorting, immunoprecipitation of BrdU-labelled DNA and molecular cytogenetics. Y.J. did the Repli-Seq libraries and the sequencing. S.G., S.E.-H. and C.-L.C. designed and performed sequencing data analyses and statistical analyses. O.B., S.G. and D.A. contributed equally to this work.

## DATA availability

All sequencing files and processed count matrices were deposited in Gene Expression Omnibus (GEO). Previously published data (accessions numbers) have been included in the Methods section where appropriate.

## Code availability

The computer codes and further processing data are available on the corresponding GitHub repositories of the team (https://github.com/CL-CHEN-Lab/).

