## supplemental figures and tables for "Unscheduled origin building in S-phase upon tight CDK1 inhibition suppresses CFS instability"

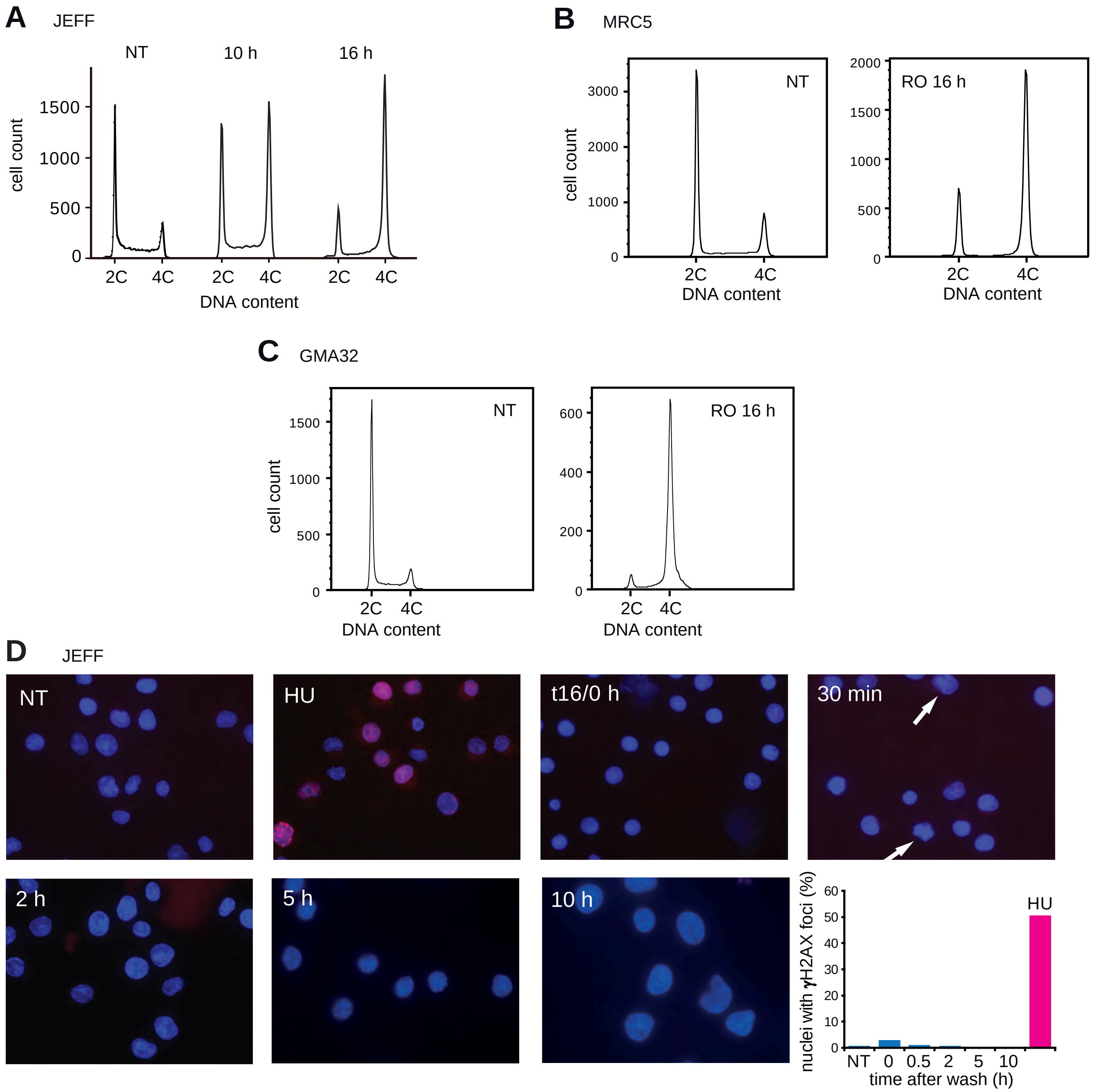

**Figure S1 A-D: RO reversibly blocks cells in G2 phase, without triggering DNA damage. A:** JEFF cells were incubated for 10 or 16 h in the presence RO (10 µM), fixed, stained with propidium iodine and analyzed by FACS. **B, C:** MRC5 human primary fibroblasts and GMA32 immortalized Chinese hamster fibroblasts were treated for 16 h with RO, and analyzed as in A. **D:** JEFF cells were treated as in figure 1A. At indicated times, samples were collected and analyzed by IF with anti-γH2AX antibodies. The percentage of nuclei with foci in each sample was determined by eye counting (histogram). Note the presence of mitotic cells (white arrows) in the 30 min panel.

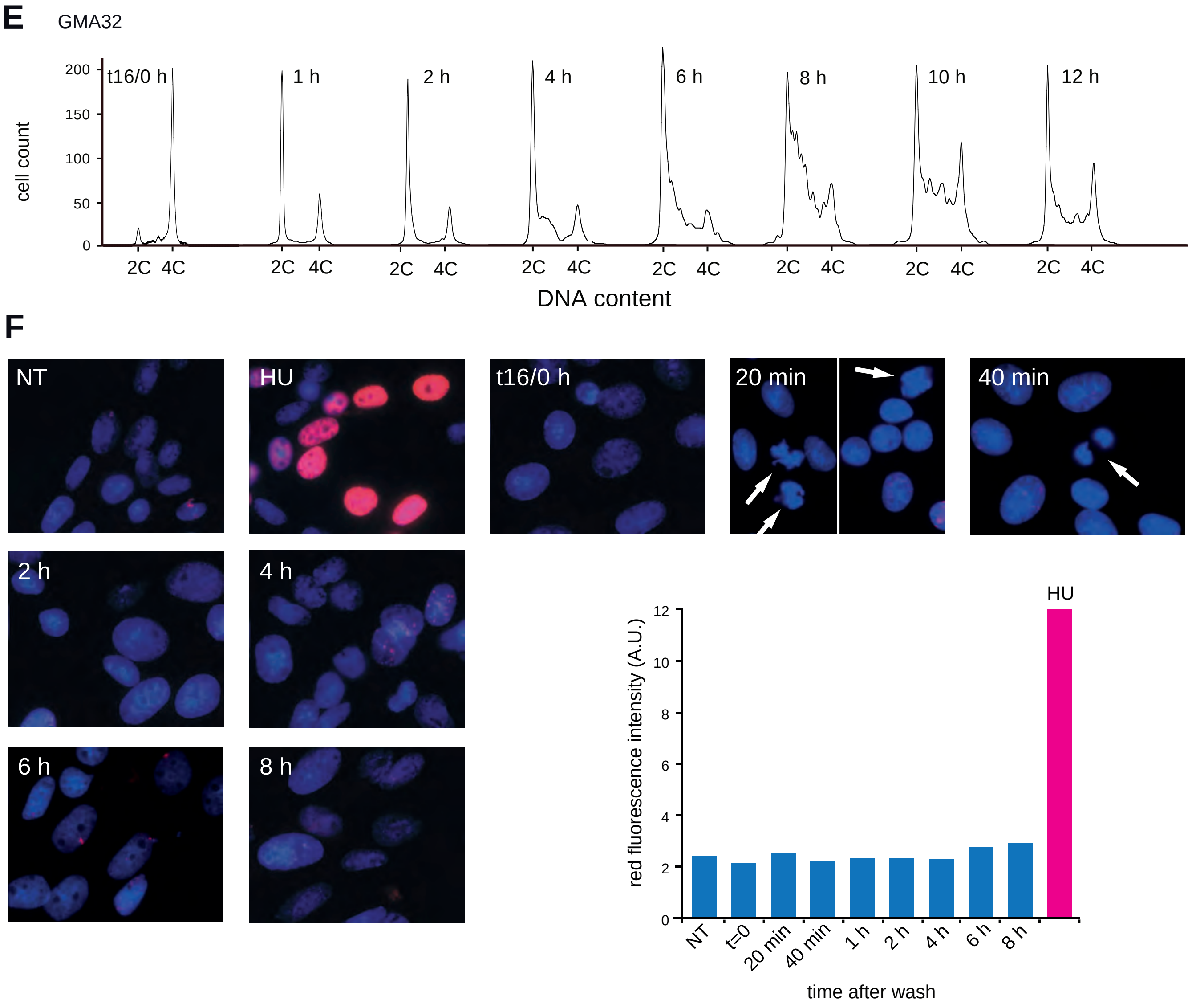

**Figure S1 E, F:** GMA32 cells were treated and analyzed at the indicated times as in figure 1A. For quantification (histogram), pictures of 20 microscopic fields for each sample were assembled and the intensity of global red fluorescence was determined with Image Gauge.

**G**

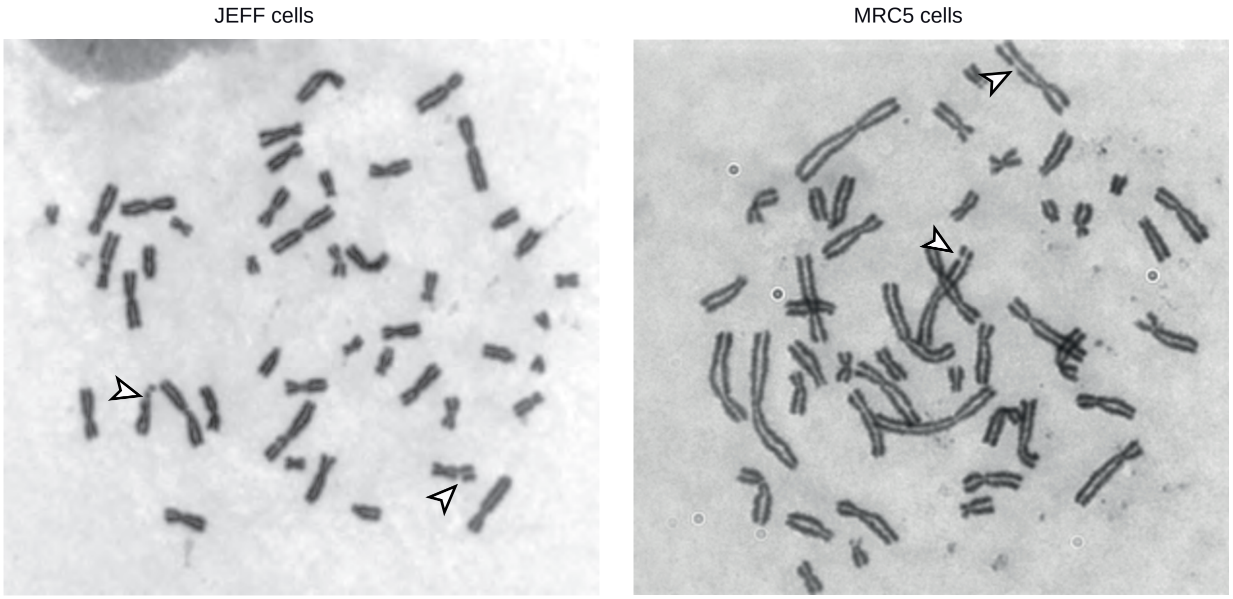

**H**

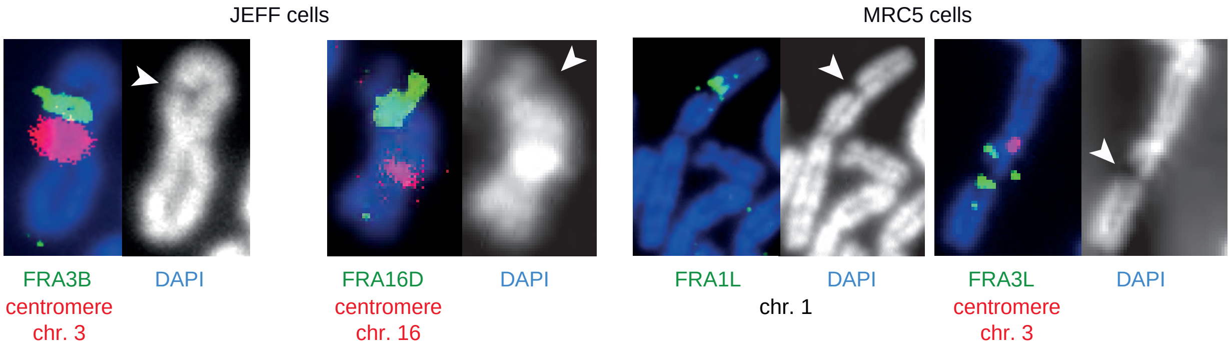

**I**

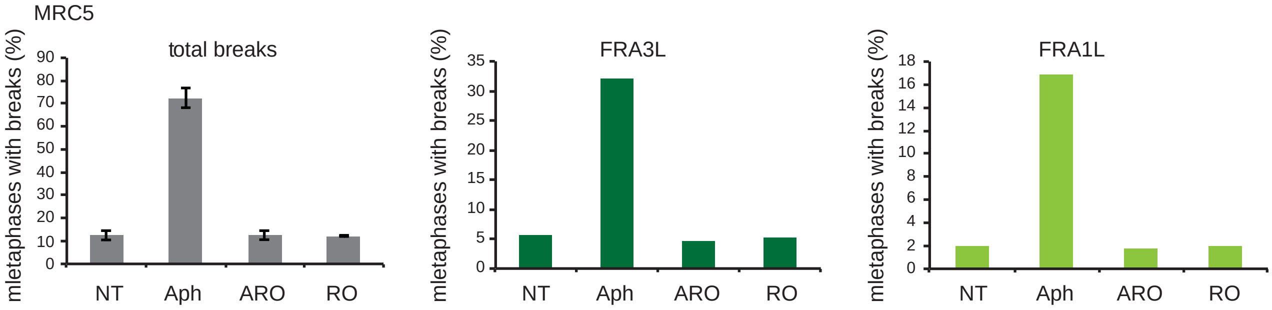

**Figure S1 G-I: Conventional and molecular cytogenetic analyses of cells treated with Aph or ARO. G:** Metaphase plates from JEFF or MRC5 cells treated with Aph 600 nM for 16 h. Chromosomes were stained with Giemsa, arrowheads point to breaks. **H:** Examples of chromosome breaks (white arrowheads) at CFSs in each cell type. Breaks at FRA3B/*FHIT* or FRA16D/*WWOX* in JEFF cells and at FRA1L/*NEGR1* or FRA3L/*LSAMP* in MRC5 cells are shown. Left panels: FISH with probes specific to fragile genes (green) in association with probes specific to the centromere of corresponding chromosomes when available (red). Right panels: Chromosomes counter stained with DAPI. Contrast was enhanced to clearly show the breaks. **I:** Total breaks and breaks at FRA3L and FRA1L were determined. Results are presented as in figures 1E.

J

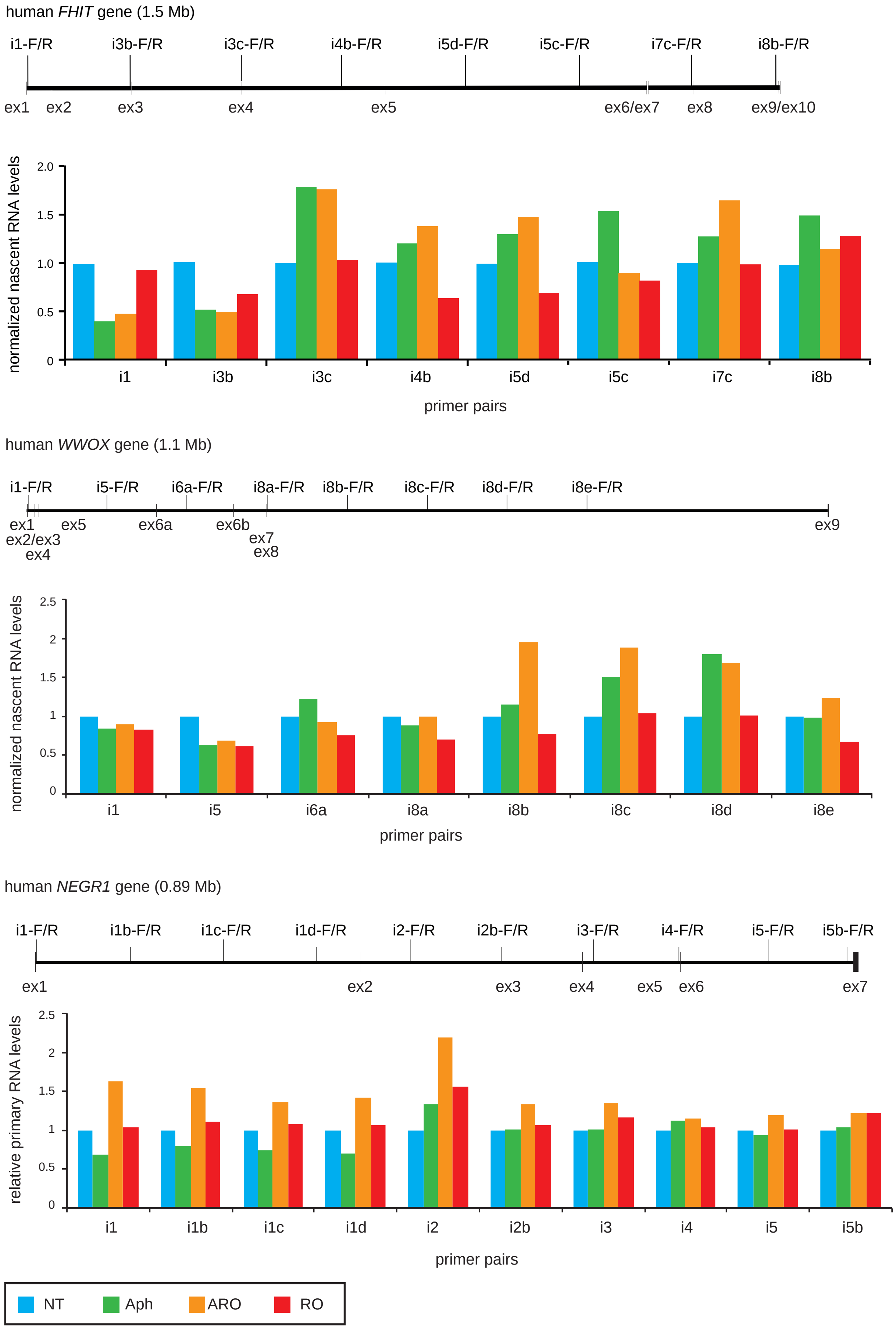

human FHIT gene (1.5 Mb)
i1-F/R
i3b-F/R
i3c-F/R
i4b-F/R
i5d-F/R
i5c-F/R
i7c-F/R
i8b-F/R
ex1
ex2
ex3
ex4
ex5
ex6/ex7
ex8
ex9/ex10
normalized nascent RNA levels
2.0
1.5
1.0
0.5
0
i1
i3b
i3c
i4b
i5d
i5c
i7c
i8b
primer pairs
human WWOX gene (1.1 Mb)
i1-F/R
i5-F/R
i6a-F/R
i8a-F/R
i8b-F/R
i8c-F/R
i8d-F/R
i8e-F/R
ex1
ex2/ex3
ex4
ex5
ex6a
ex6b
ex7
ex8
ex9
normalized nascent RNA levels
2.5
2
1.5
1
0.5
0
i1
i5
i6a
i8a
i8b
i8c
i8d
i8e
primer pairs
human NEGR1 gene (0.89 Mb)
i1-F/R
i1b-F/R
i1c-F/R
i1d-F/R
i2-F/R
i2b-F/R
i3-F/R
i4-F/R
i5-F/R
i5b-F/R
ex1
ex2
ex3
ex4
ex5
ex6
ex7
relative primary RNA levels
2.5
2
1.5
1
0.5
0
i1
i1b
i1c
i1d
i2
i2b
i3
i4
i5
i5b
primer pairs
NT
Aph
ARO
RO

**K**

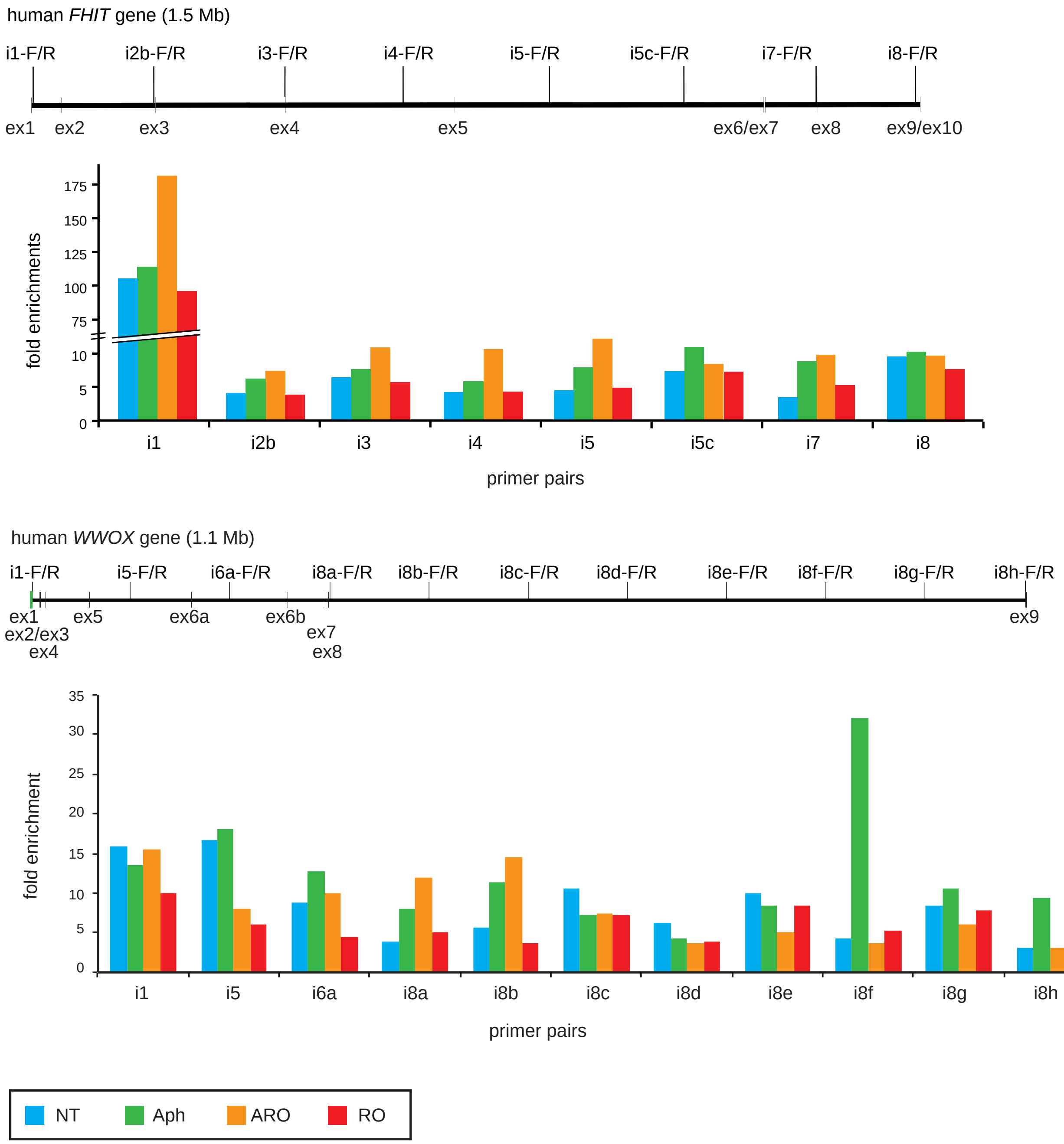

**Figure S1 J, K: Analysis of the transcription level of large fragile genes in cells treated with Aph, RO or ARO. J:** Comparison of nascent RNA levels along the *FHIT* and *WWOX* genes in JEFF cells, and of primary RNA levels along the NEGR1 gene in MRC5 cells. Untreated (NT) cells and cells grown for 16 h in either medium (as in figure 1C) were pulse-labeled with 5-ethynyl-uridine. Nascent RNA were purified by affinity chromatography and quantified by RT-qPCR using *FHIT*, *WWOX* or *NEGR1* intronic primers whose positions are indicated on the gene maps together with exon (ex) position. Results are presented relative to the level in NT. **K:** JEFF cells were treated as in J. Fixed chromatin was prepared and the density of RNA polymerase II along the *FHIT* and *WWOX* genes was determined by immuno-precipitation with antibodies anti-RNA polymerase II. Quantification was done by qPCR using the same intronic primers as above. Results are presented as fold enrichment relative to the enrichment obtained with immunoglobulins from a normal rabbit. Note the high level of polymerase accumulation on the *FHIT* promoter detected by the i1 primers whose positions are close to the small exon 1.

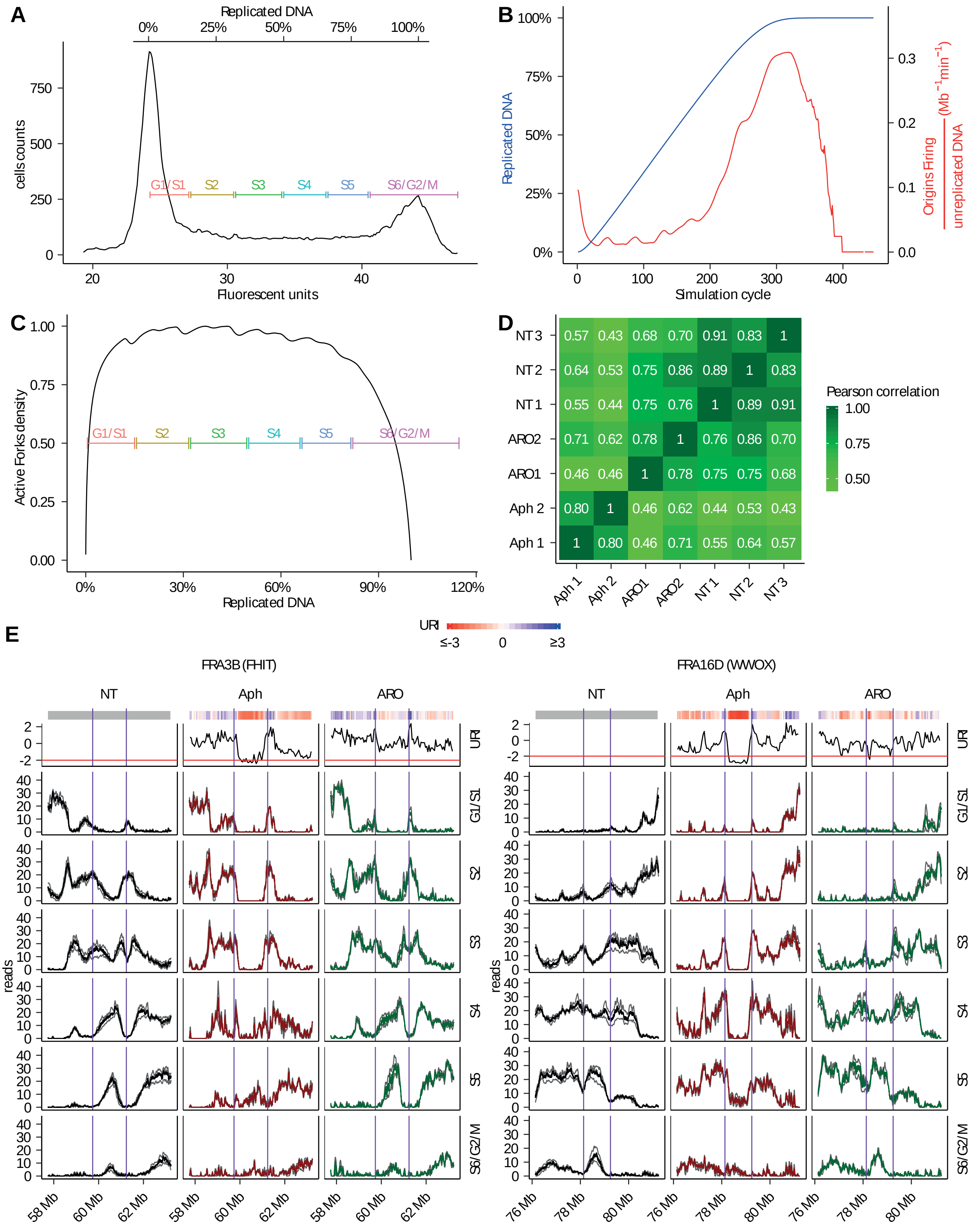
A
Replicated DNA
0%
25%
50%
75%
100%
cells counts
750
500
250
0
G1/S1
S2
S3
S4
S5
S6/G2/M
20
30
40
Fluorescent units
B
100%
75%
50%
25%
0%
Replicated DNA
0.3
0.2
0.1
0.0
Origins Firing
unreplicated DNA
(Mb⁻¹min⁻¹)
0
100
200
300
400
Simulation cycle
C
1.00
0.75
0.50
0.25
0.00
Active Forks density
G1/S1
S2
S3
S4
S5
S6/G2/M
0%
30%
60%
90%
120%
Replicated DNA
D
NT3 0.57 0.43 0.68 0.70 0.91 0.83 1
NT2 0.64 0.53 0.75 0.86 0.89 1 0.83
NT1 0.55 0.44 0.75 0.76 1 0.89 0.91
ARO2 0.71 0.62 0.78 1 0.76 0.86 0.70
ARO1 0.46 0.46 1 0.78 0.75 0.75 0.68
Aph 2 0.80 1 0.46 0.62 0.44 0.53 0.43
Aph 1 1 0.80 0.46 0.71 0.55 0.64 0.57
Aph 1
Aph 2
ARO1
ARO2
NT1
NT2
NT3
Pearson correlation
1.00
0.75
0.50
E
URI
≤-3
0
≥3
FRA3B (FHIT)
FRA16D (WWOX)
NT
Aph
ARO
URI
G1/S1
S2
S3
S4
S5
S6/G2/M
reads
58 Mb
60 Mb
62 Mb
76 Mb
78 Mb
80 Mb

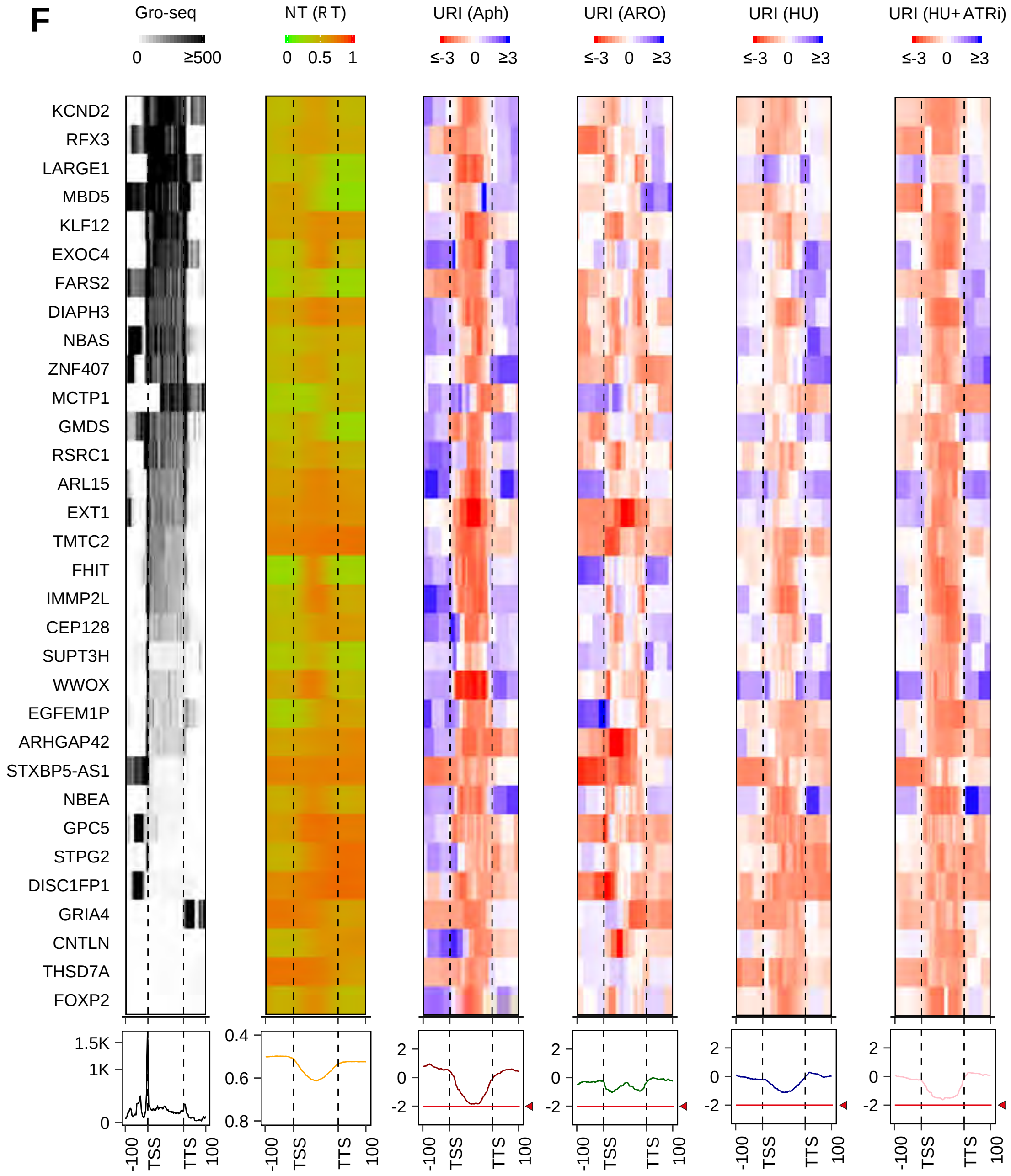

**Figure S2: Repli-Seq data analyses. A:** Representative FACS profile showing the gate position for cell sorting. Fluorescent units are reported on the x axis and the cell counts on the y axis. The percentage of replicated DNA (based on the G1 and G2 peak position) is shown on the top. **B:** Simulation of S phase progression (based on [59]). Blue curve: percentage of replicated DNA shown as a function of simulation cycle. Red curve: typical bell shaped curve given by the ratio between origin firing and un-replicated DNA as a function of simulation cycle. **C:** Density of active forks in function of the percentage of replicated DNA obtained in the simulation. The relative quantity of newly replicated DNA inside each fraction has then been used to normalize the Repli-Seq data. **D:** Heat-map showing Pearson correlation coefficient between individual biological replicate of NT, Aph and ARO samples. **E:** Repli-Seq profiles of FRA3B and FRA16D showing the profiles of individual replicates in each condition in grey, and the average profiles in blue for NT, red for Aph and green for ARO (as in figure 2B). **F:** Heatmaps and average profiles of transcription levels (GRO-seq), replication timing in non-treated cell (RT (NT)) and URI in Aph, ARO or HU (as in figure 3) over large genes hosting SDRs. Genes are ordered based on decreasing GRO-seq signal as in figure 3 and the name of each gene is reported on the left. Red lines and arrowheads on URI average profiles: as in Figure 2B. Note that the average profile does not reach the -2 threshold for genes containing SDRs because the SDWs are not located at the same place in the different genes.

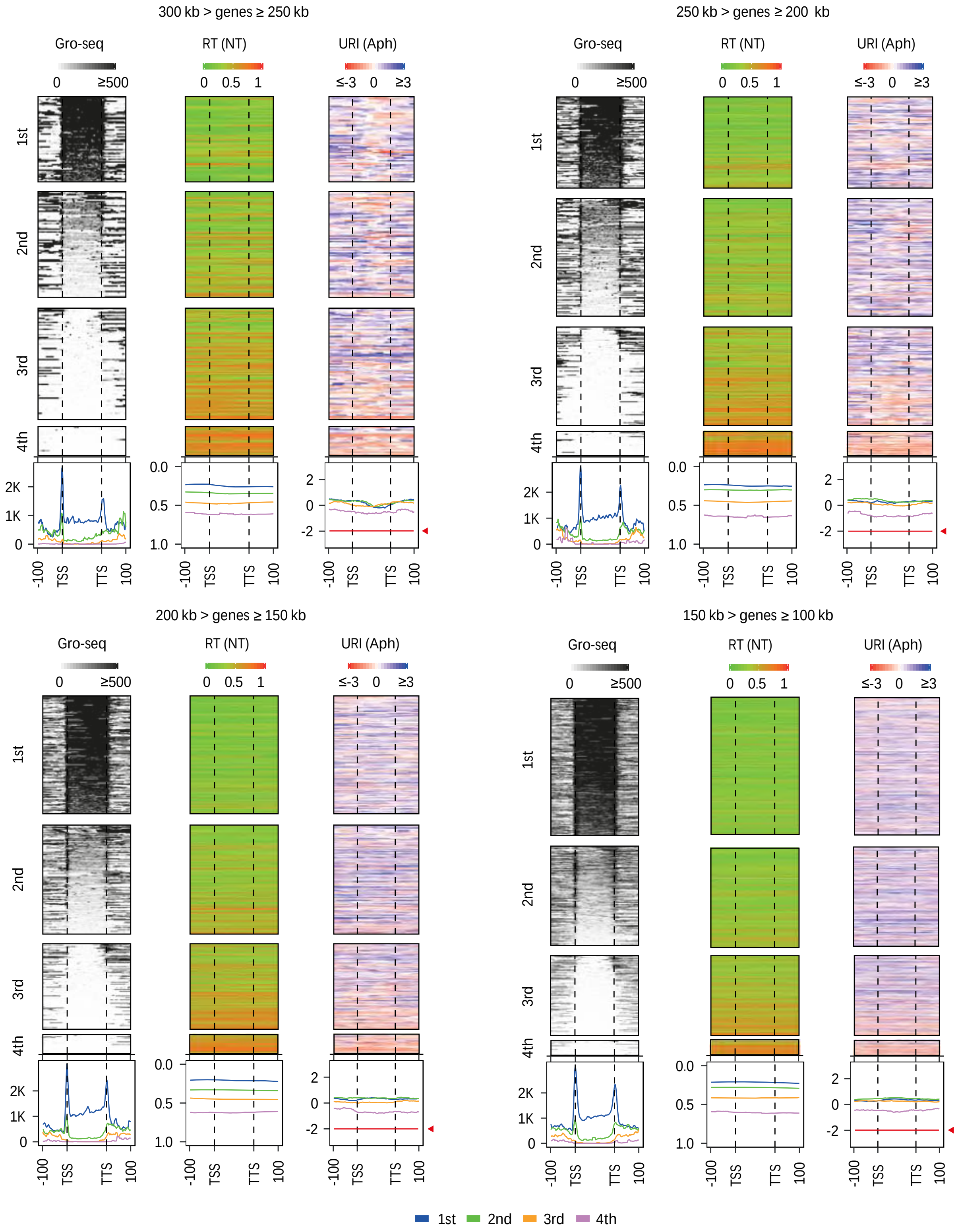

**Figure S3: Analysis of Gro-seq, RT and URIs across genes 100- to 300-kb long.** Heat-maps and mean profiles of GRO-seq, RT and Aph-induced URIs (as in figures 3 and S2F) are shown for genes belonging to the indicated classes of size. Smaller genes were not studied because the current resolution of the Repli-Seq technique is ≈ 50 kb. As in figure 2B, S2 and 3, GRO-seq values over 500 counts and URI values below -3 or above 3 have been saturated for visualization purposes in all the heatmaps. Red lines and arrowheads: as in figure 2B.

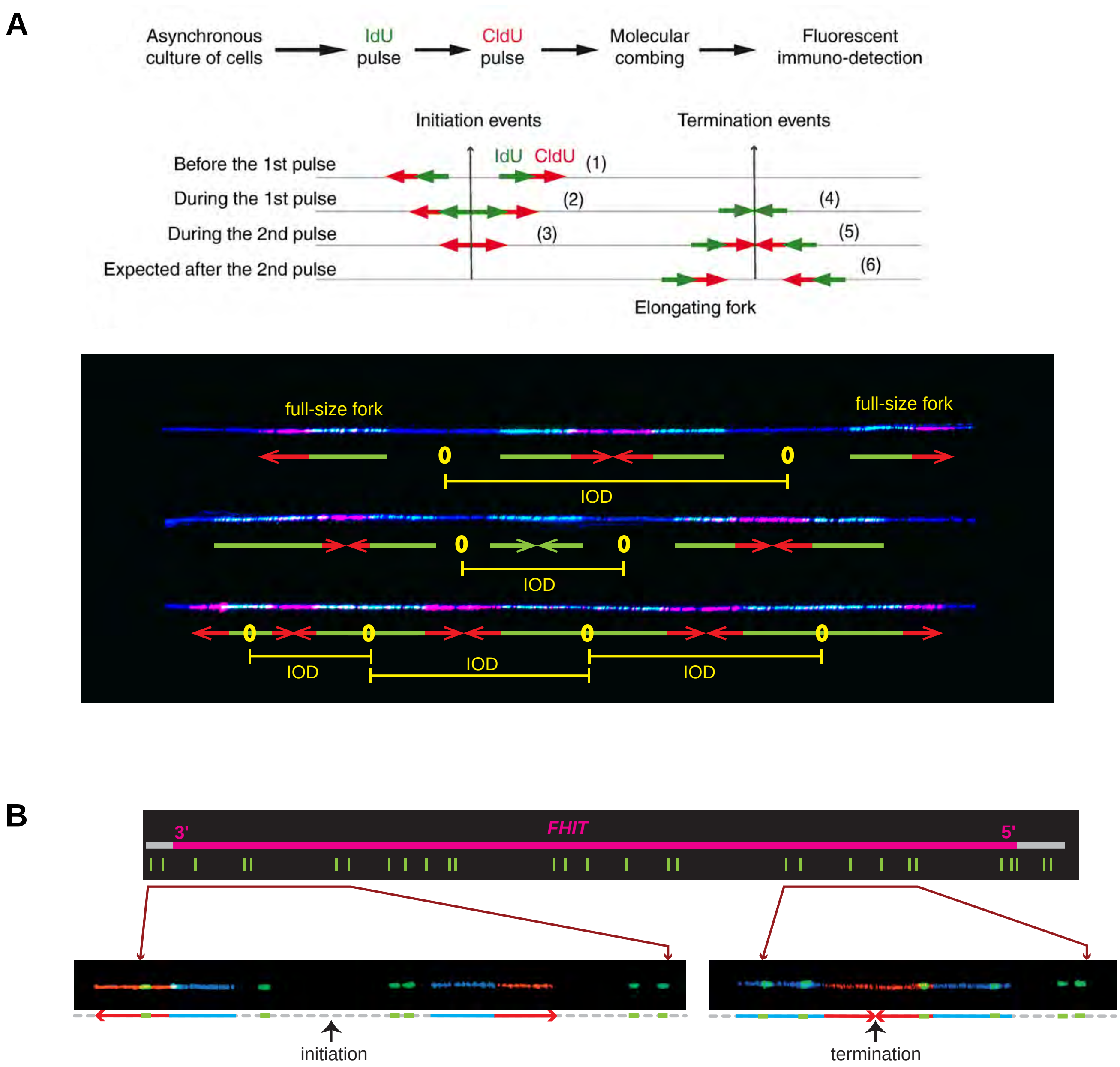

**Figure S4: Principles and examples of replication dynamics analyses. A:** Upper panel: Scheme of the protocol used for replication dynamics analyses at the whole genome level in exponentially growing cells. Successive IdU and CldU pulses enable recognition of the direction of fork progression, therefore of initiation and termination events. Green and red arrows represent neo-synthesized DNA labeled with IdU or CldU, respectively. The labeling pattern depends on whether the considered event had occurred before the first pulse, during the first pulse, during the second pulse or after the second pulse. Six different patterns of initiations and terminations are expected, numbered 1–6 (adapted from [60]). Lower panel: Examples of DNA fibers (DNA counterstaining in blue) harboring initiation events with schematic representation shown below each fiber. Arrowheads indicate the direction of fork movement (as in A); O: origin; IOD: inter-origin distance (distance separating two adjacent initiation events in clusters of origins firing more or less at the same time). Examples of full-size forks used for speed determination are shown. **B:** Upper panel: FISH Morse code used for identification of a region 1.6 Mb-long encompassing the *FHIT* gene. It comprises 29 probes (green bars) organized in five motifs. Lower panels: Examples of DNA fibers bearing Morse code motifs (green) and replication signals (newly synthesized DNA labeled as above except that IdU is revealed in blue). A schematic representation of the replication tracks and FISH signals is shown below each fiber. Arrowheads indicate the direction of fork progression. Black arrows indicate the barycenter of forks or the estimated positions of initiation and termination events (adapted from [48]).

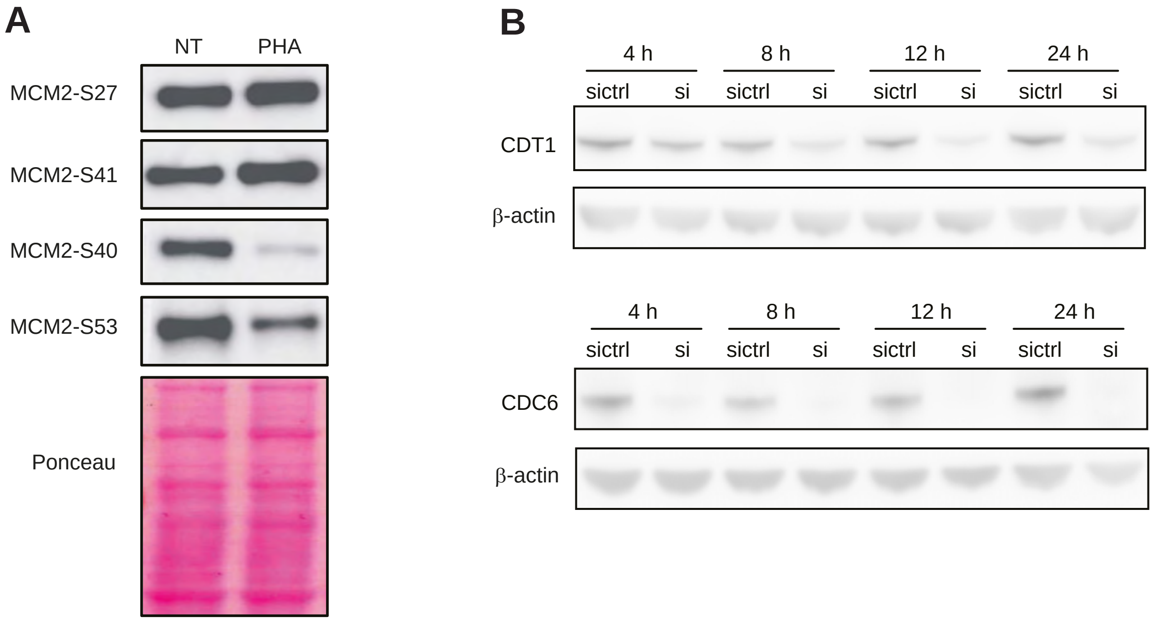

**Figure S5 A, B: Stability of CFSs in ARO treated cells relies on extra initiation events, CDC6 and CDT1. A:** JEFF cells were untreated (NT) or treated with 6 µM PHA-767491 (PHA) for 16 h. Total protein extracts were analyzed by Western blotting with antibodies specific to the phosphorylated forms of MCM2 on ser27, ser41, ser40 or ser53. Ponceau staining of the membrane was used as a loading control. **B:** JEFF cells were transfected with siRNAs targeting human CDT1 or CDC6 mRNAs (siCDC6, siCDT1), or with a control siRNA with no known target (sictrl). At indicated times after transfection, total protein extracts were prepared to assess the level of each factor by Western blotting. Beta-actin was used as a loading control.

C

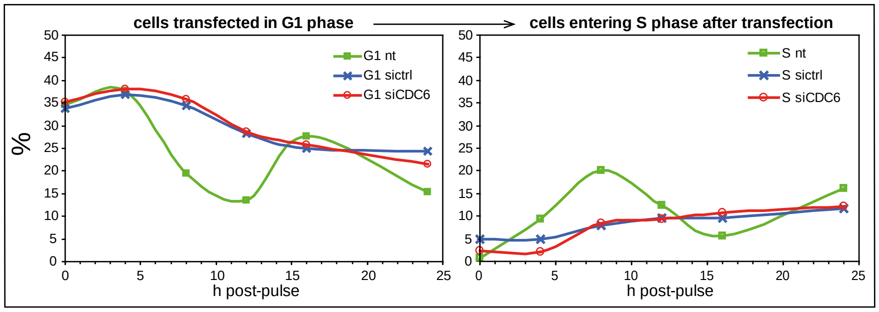

D

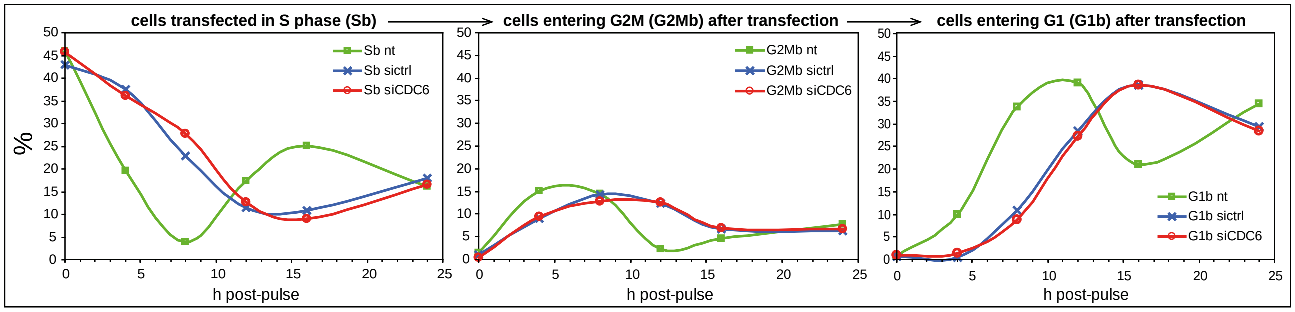

E

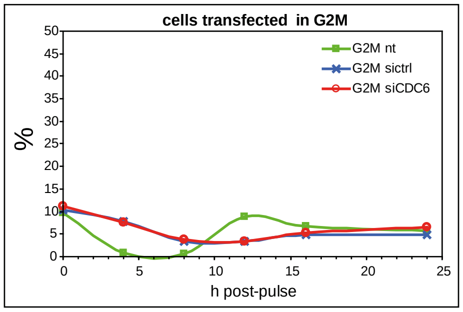

**Figure S5: C-H: Resilience of cells transfected at different stages of the cell cycle.** JEFF cells were pulse labeled for 2 h with BrdU just after transfection with the indicated siRNAs and cell cycle progression was followed thereafter by FACS analysis. Cells transfected with sictrl and siCDC6 were analyzed 4 h, 8 h, 12 h, 16 h, and 24 h post transfection in parallel of non-transfected cells (nt) for comparison (F, G, H). Fractions of the cells in each phase of the cell cycle were quantified (percentages indicated in red polygons of each panel). Sb, BrdU-labeled cells (FITC fluorescence). **C:** Non transfected cells that were in G1 at the time of BrdU pulse (unlabeled G1) exit this phase, enter S phase (unlabeled S) and are back in G1 by 16 h with kinetics fully compatible with the known parameters of JEFF cell cycle (G1 : 6 h; S : 8 h; G2M : 2 h). In striking contrast, cells transfected in G1 exit this phase and enter S phase extremely slowly. By 24 h post-transfection, only a small fraction of them have progressed past these phases. **D, E:** Non-transfected cells that were in S (labeled S) (E) or G2M (unlabeled G2/M) (E) at the time of BrdU pulse progress as expected in the cycle. In contrast, cells transfected in S (D) or G2/M (E) phase exit these phases with a 6-h delay. This delay is still visible when cells transfected in S phase progress in G2M (labeled G2/M) and then in G1 (labeled G1) (D).

**F**

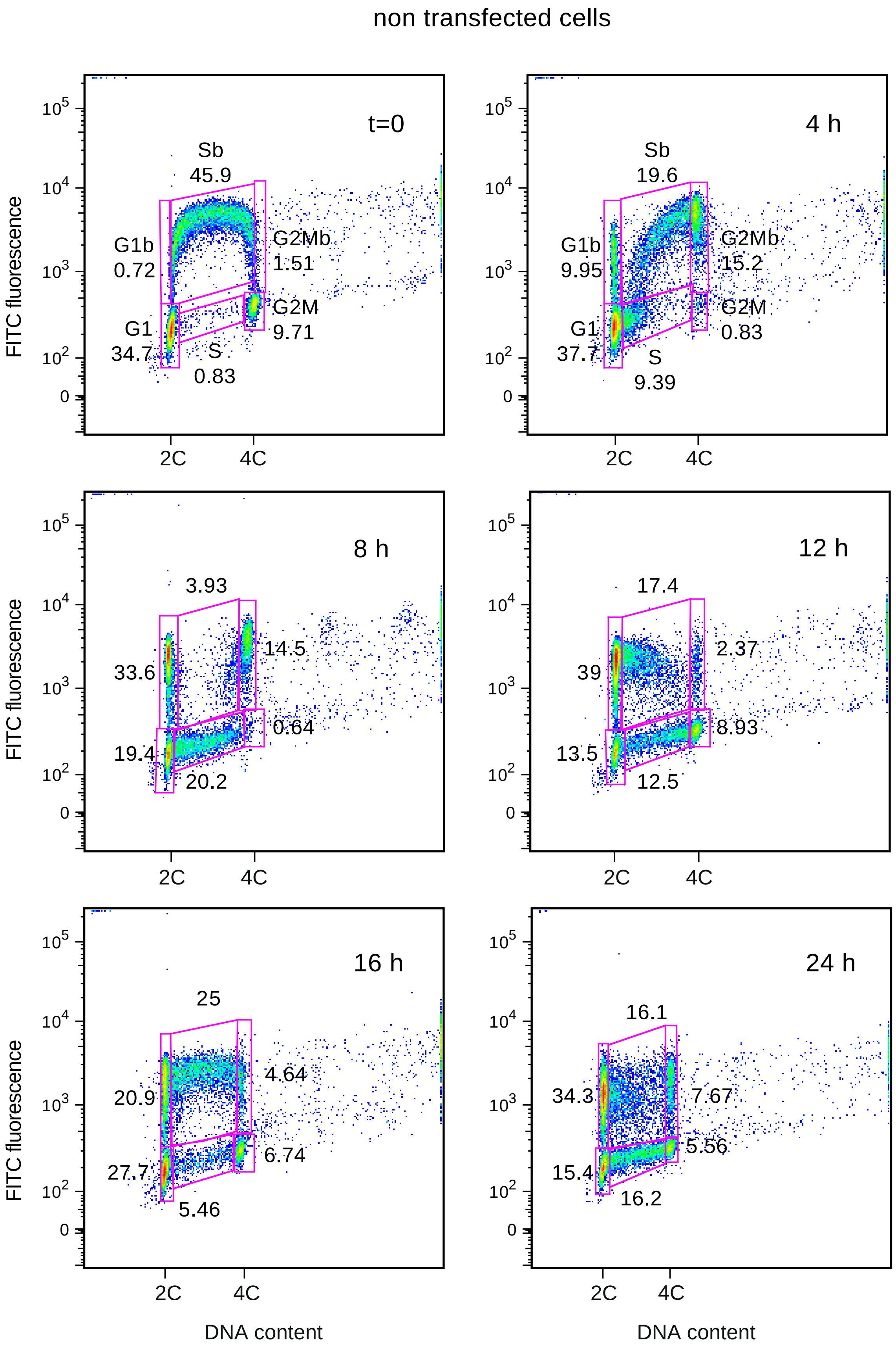

non transfected cells
t=0
Sb
45.9
G1b
0.72
G2Mb
1.51
G1
34.7
G2M
9.71
S
0.83
4 h
Sb
19.6
G1b
9.95
G2Mb
15.2
G1
37.7
G2M
0.83
S
9.39
8 h
3.93
33.6
14.5
0.64
19.4
20.2
12 h
17.4
39
2.37
8.93
13.5
12.5
16 h
25
20.9
4.64
27.7
6.74
5.46
24 h
16.1
34.3
7.67
5.56
15.4
16.2
FITC fluorescence
DNA content
2C
4C

**G**

transfection with sictrl siRNA

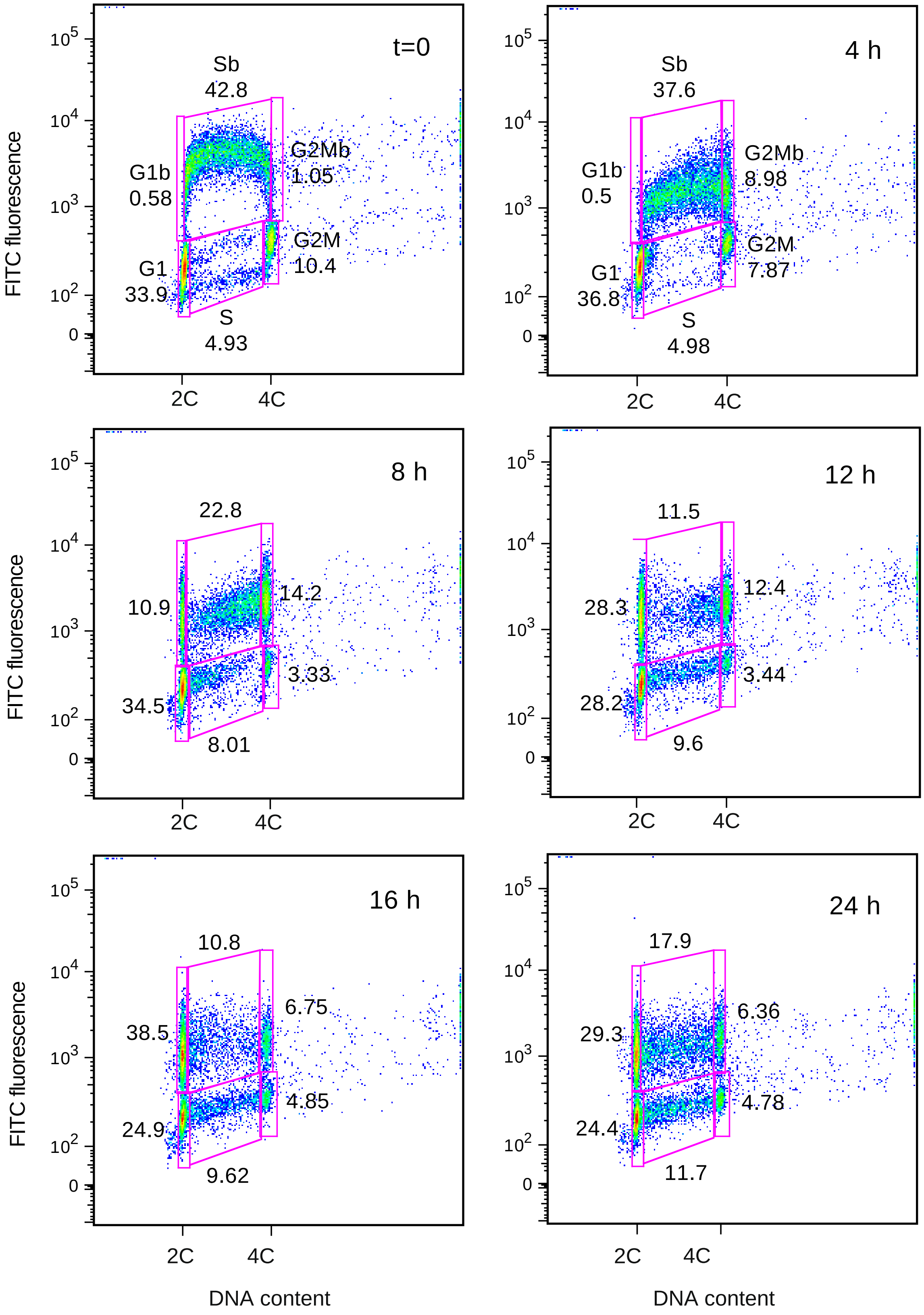

**H**

transfection with siCDC6 siRNA

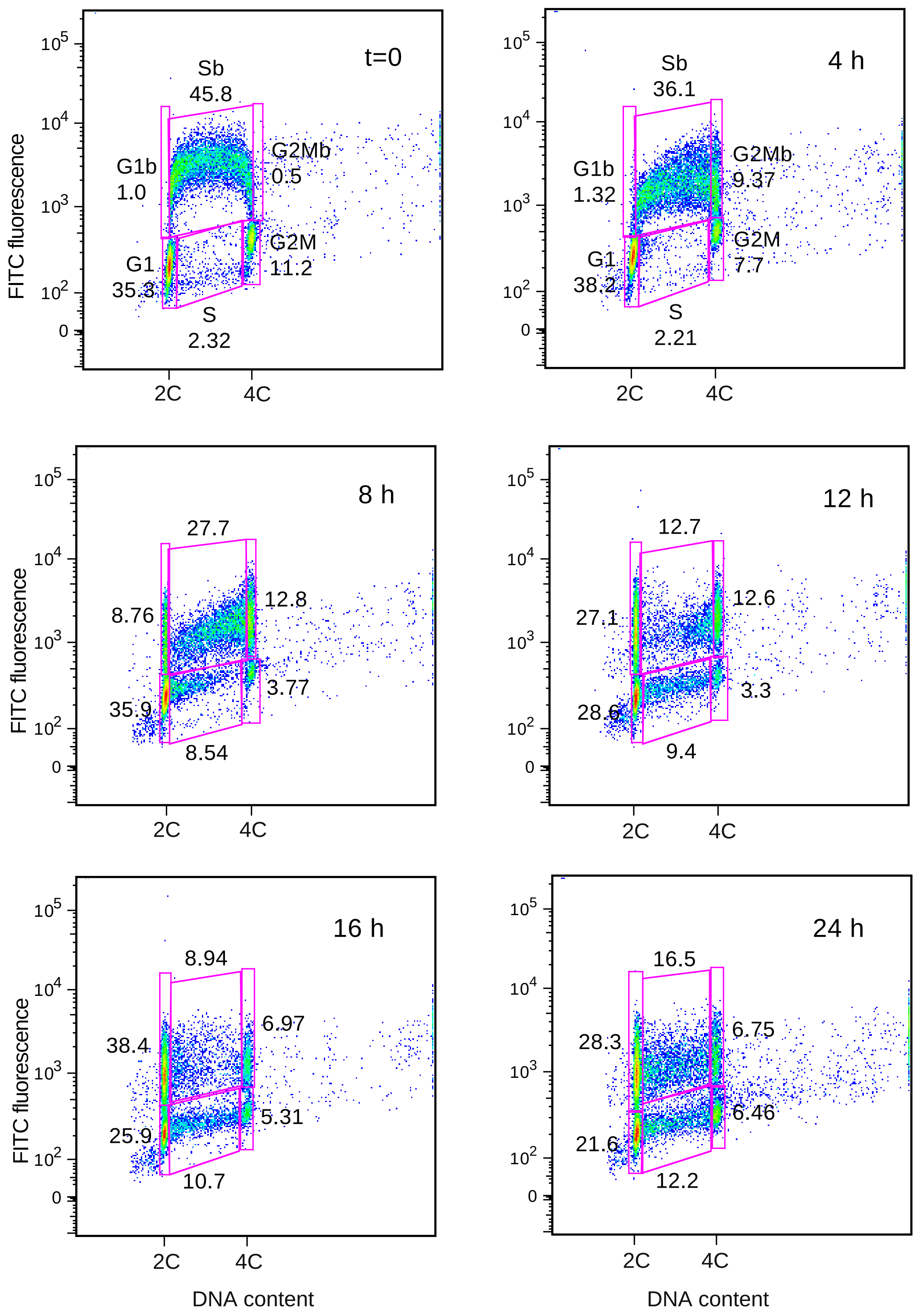

### Supplemental Table I

| antibody | supplier | reference | working dilution |
|---|---|---|---|
| rabbit monoclonal anti-CDC6 | Cell Signaling | 3387 | 1/1000 |
| rabbit monoclonal anti-CDT1 | Cell Signaling | 8064 | 1/1000 |
| mouse monoclonal anti-MCM7 | Santa Cruz | sc-56324 | 1/1000 |
| rabbit monoclonal anti-MCM2 phospho-S27 | Abcam | ab109459 | 1/10 000 |
| rabbit monoclonal anti-MCM2 phospho-S40 | Abcam | ab133243 | 1/1000 |
| rabbit monoclonal anti-MCM2 phospho-S41 | Abcam | ab109270 | 1/50 000 |
| rabbit monoclonal anti-MCM2 phospho-S53 | Abcam | ab109133 | 1/10 000 |
| mouse monoclonal anti-$\beta$-actine-HRP | Santa Cruz | sc-47778 HRP | 1/20000 |
| rabbit polyclonal anti-CHK1 phospho-Ser317 | Cell Signaling | 2344 | 1/500 |
| rabbit polyclonal anti-p53 phospho-Ser15 | Cell Signaling | 9284 | 1/500 |
| rabbit polyclonal anti-RNA polymerase II CTD repeat phospho-S5 | Abcam | ab5131 | n. a. |
| rabbit polyclonal anti-histone H3 | Abcam | ab1791 | 1/1000 |
| mouse monoclonal anti-MPM2 | Millipore | 05-368 | 1/500 |
| rat monoclonal anti-BrdU | AbD Serotec | MCA2060 | 1/100 |
| mouse monoclonal anti-phospho-histone H2AX | Merck | 05-636 | 1/400 (FACS) or 1/5000 (IF) |
| rabbit monoclonal anti-gamma-H2AX phospho-Ser139 | Abcam | ab81299 | 1/1000 |
| rabbit polyclonal anti-gamma-H2AX phospho-Ser139 | Abcam | 2893 | 1/1000 |
| rabbit polyclonal immunoglobulins - Isotype Control | Abcam | ab37415-5 | n. a. |
| goat polyclonal anti-mouse immunoglobulins-HRP | Dako | P0447 | 1/5000 |
| goat polyclonal anti-rabbit immunoglobulins-HRP | Dako | P0448 | 1/5000 |
| chicken polyclonal anti-rat immunoglobulins-Alexa 488 | Invitrogen | A-21470 | 1/50 |
| goat polyclonal anti-mouse immunoglobulins-Alexa 488 | Invitrogen | A11029 | 1/200 |
| goat polyclonal anti-rabbit immunoglobulins-Alexa 594 | Invitrogen | A11012 | 1/400 |
| goat polyclonal anti-mouse immunoglobulins-Alexa 594 | Invitrogen | A11032 | 1/1000 |

n. a. not applicable

**Supplemental Table II**

| gene | Forward primer | Reverse primer |
|---|---|---|
| FHITi1 | TGTGCTGGGACCAATAGAAA | AACATCAGGTGCGAGTAGGG |
| FHITi2b | TTGGCTAGGAAACGGAATTCC | GCAGGGCAGGCATTGC |
| FHITi3 | TTGGGGGAACAGAATTCAAC | CATGCTGCCTACCTTCTGGT |
| FHITi3b | CATCCAGTGTCCAAGATGCATAC | CCCTGCCTCCCCTTCCT |
| FHITi3c | AGTGCATTGTCATTGAGTGATTTATG | GATAACCCCAAAGTGGACAAGTG |
| FHITi4 | TGGCATATTGGACAGGGGAGGT | CTAGTGCGGGACCTGGCACA |
| FHITi4b | GGCTTCGTGGGAGAAAAGG | AGATGCTGCTGTTGACAGTCGTA |
| FHITi5 | TGGGCAACACACATCTGGAACA | TCACTGGGTCAGTGCCTGCTC |
| FHITi5c | ACCGTAGGGAACCTCCTCTTG | GCTGCTCTTTGACCTTGAACAGT |
| FHITi5d | TCTCTCCTTTCAGGCTCCAAAT | AAGAAACAGGCCAAGTTAATGCA |
| FHITi7c | CAGCTCCCTTTCCTAGTCTGGAT | GGAAGCATTGATTGCAGATGAG |
| FHITi8b | TGGCGTGCACTTTCCTCTAA | CTGGTTTCCTTCTGATGCAACA |
| WWOXi1 | TCATTCAAGACACACACAGGTATTATG | TGGCTCTTATTTGCCTCGGTAT |
| WWOXi5 | AGAGGGCTAATGACACACTGCTTA | AAAAACTTTAAACAAGCCCCAAA |
| WWOXi6a | GGCGTCAGCACACACATCAC | AGAGCTGTGCTGGAAACACTAACA |
| WWOXi8a | CGCAAACACCTTGGGTCCTA | CTCATGCCCGCAAGAGGTA |
| WWOXi8b | CCGCAGGGCCTGATAGG | CAACCGTCCTTGATACAAATAAACA |
| WWOXi8c | CGGTACTTGCTTTGGCAGAAC | GATGCAGGCCAGGTGAGAA |
| WWOXi8d | CTGGGATGGTGGAGCCTAAG | TTTCTGCCTCCGCAATGC |
| WWOXi8e | AGGGTAAGATCTGCTGTCTTGTGA | CAAATTAAAGACACTGGGAGAGAGAA |
| WWOXi8f | CCTCTATGCTCCCTCATGTTTCA | AGGGCCCTGTACAGAAACAAAG |
| WWOXi8g | CCCTGGTACTGATGGTTGCA | GAAGTTTGCTCAACACACAGGAA |
| WWOXi8h | TGCCAAGATCCAGCTGAAACT | TTCGACTGCCTGGGCATT |
| NEGR1-i1 | CCCGGCCCGTTTATGC | TGGCGTTCCGGCAAAGT |
| NEGR1-i1b | TCATTTCGCCGGTTGATACC | ATGGTGACTGTGGTGCTGATG |
| NEGR1-i1c | CCAAGGGAAAGACTATTAAGGATCTC | AACAGCAAGGACTGGATTAAGCA |
| NEGR1-i1d | CATCAAGATGCTGTGTCCACTGT | CCTCACCATCTCCTGTCCCTAA |
| NEGR1-i2 | CGTGGGCAGTTCCCATTAAT | TTCATCCTGTTTGGGCATTTT |
| NEGR1-i2b | AACCTTTCTCCCATCCCAGAA | AGTTTCCCACACCCAGCTAATC |
| NEGR1-i3 | GGTGAATAGAAGAATGGAGAGAAAAATAA | TTGCCTCTCCACGTGTGTACTC |
| NEGR1-i4 | AAGCAGAAGTAGGTACAACATGCATT | ACCACCAGAGGCCCTCACT |
| NEGR1-i5 | GCCTTTGAAGCGACTTTGGA | CTGCCTGCCAGGTCTCTTG |
| NEGR1-i5b | ACCCATTCCAAACACTCCAATAA | ACTTGGCTAACAGTCATGTTGAATTT |
| cyclophilin B (PPIB) | GTGAGCGCTTCCCCGATGAG | TGCCAAACACCACATGCTTGC |
